# Analysis of ribosomes from WT and FMR1 KO human embryonic stem cells

**DOI:** 10.1101/2024.11.18.624098

**Authors:** Tripti Kharbanda, Michelle Ninochka D’Souza, Dasaradhi Palakodeti, Ravi S Muddashetty, Kutti R. Vinothkumar

## Abstract

Fragile X Messenger Ribonucleoprotein 1 (FMRP) is a multifunctional, multidomain RNA-binding protein whose loss causes Fragile X syndrome. It is also known to associate with ribosomes and modulate translation. In human embryonic stem cells (hESCs), knockout (KO) of the FMR1 gene results in significantly increased protein translation rates and alterations in the 2’-O-methylation patterns of rRNA. To understand the structural underpinnings of the process, we performed structural analysis of ribosomes isolated from both the wild-type (WT) and the FMR1 KO hESCs, which revealed populations of ribosomes with distinct protein factors and tRNA in the FMR1 KO ribosome preparations. The bound protein factors are hallmarks of dormant ribosomes, which are currently not observed in the WT hESCs, and are characterized by the binding of SERPINE-mRNA binding protein 1 in the mRNA tunnel and eukaryotic elongation factor 2 near the A-site. The observation of elevated protein translation in FMR1 KO cells, alongside a subpopulation of dormant ribosomes, further substantiates the role of FMRP in translational regulation. Additionally, we analysed the 2’-O-methylation patterns in the 28S rRNA, in both the WT and the FMR1 KO hESCs, identifying a few potential differentially methylated sites. Thus, these observations provide insights into the mechanisms of ribosome dormancy in the absence of FMRP and in understanding the role of rRNA methylation in translational regulation.

## Introduction

Fragile X Messenger Ribonucleoprotein 1 (FMRP, previously called Fragile X mental retardation protein) is a multifunctional protein implicated in various functions including translational regulation, maintenance of synaptic plasticity, RNA transport, and maintaining ion channel stability (Davis & Broadie, 2017; Goering *et al*., 2020; Lai *et al*., 2020; Deng *et al*., 2013; Brown *et al*., 2010; Bhakar *et al*., 2012). The loss of this protein results in Fragile X syndrome (FXS), a common inherited neurodevelopmental disorder leading to intellectual disability (Santoro *et al*., 2012). FXS is caused by the expansion of a trinucleotide CGG repeat in the 5’ untranslated region of the FMR1 gene, which results in DNA hypermethylation and subsequent transcriptional silencing, leading to the absence of the FMR1 gene product (M H Verkerk *et al*., 1991). FMRP has been shown to be present in both the nucleus and the cytoplasm and has the consensus motifs for both nuclear localization and nuclear export signals (Eberhart *et al*., 1996). Structurally, FMRP is a multidomain protein with multiple RNA-binding regions, namely, 3 K-homology (KH0, KH1, and KH2) domains and an Arginine-glycine-glycine box (RGG) at the C-terminus. KH0 is present towards the N-terminus with 2 Agenet/Tudor domains known to interact with other proteins (Myrick *et al*., 2015).

Among its physiological functions, translational regulation is one of the most well-studied roles of FMRP. It is proposed that FMRP regulates specific transcripts by binding to the UTR regions of mRNA or through mediation by small RNAs (Taha & Ahmadian, 2024; Darnell & Klann, 2013; Richter & Zhao, 2021; Darnell *et al*., 2011; Bhakar *et al*., 2012). A direct mode of FMRP-mediated translation regulation is by binding to ribosomes, proposed to be near the ribosomal protein L5 (Chen *et al*., 2014; Athar & Joseph, 2020; D’Souza *et al*., 2022). The RNA binding activity of FMRP is critical for its interaction with the polysomes and also for regulating the translation of specific mRNAs (Darnell *et al*., 2011; Chen *et al*., 2014). Another less-explored modality of FMRP-mediated translation regulation is via its interaction with C/D Box snoRNAs in the nucleus, which target and assist in methylation of the residues in the rRNA of ribosomes (D’Souza *et al*., 2018). This further illustrates the growing regulatory role of FMRP beyond its implication in FXS for which it is typically known.

Methylation is one of the common chemical modifications found in RNAs, and among these, 2’-O-methylation of the ribose along with pseudouridinylation is the most abundant (Erales *et al*., 2017; Natchiar *et al*., 2018, 2017; Krogh *et al*., 2016). For instance, 2 dimethyl adenines (m^6^_2_A 1850 and m^6^_2_A 1851) in 18S rRNA are the most conserved base modifications across the kingdoms of life and they facilitate the subunit interactions between A- and P-sites of the ribosome (Natchiar *et al*., 2018; Poldermans *et al*., 1980; Zorbas *et al*., 2015). The methylation of the 2’ hydroxyl of the ribose is thought to provide greater stability, and the lack of hydroxyl results in altered hydrogen bonding due to which nucleophilic character and protein binding might be affected (Ayadi *et al*., 2019). Over 100 sites are methylated, and while some of these sites have established functional relevance, most remain elusive (Natchiar *et al*., 2018; Krogh *et al*., 2016; Wiechert *et al*., 2025; Faille *et al*., 2023; Holvec *et al*., 2024). Some of these modifications have been shown to be positioned close to the functional centres of the ribosomes and are thought to play an important role in translational regulation (Decatur & Fournier, 2002; Xue & Barna, 2012). It is proposed that these modifications help to maintain the structure of rRNA and fine-tune the translational regulation by creating heterogeneous pools of ribosomes that can be used under specific cellular needs and when these modifications are disrupted, it can lead to pathologies (Decatur & Fournier, 2002; Erales *et al*., 2017; Sornjai *et al*., 2017).

To understand the role of FMRP in methylation, Ribo-methylation sequencing (RMS) was performed in both the wild-type and FMR1 knockout Shef4 hESC lines, revealing differences in the rRNA 2’-O-methylation of both 28S and 18S rRNA between the WT and the FMR1 KO ribosomes (D’Souza *et al*., 2018), indicating that FMRP can bring about differential methylations in rRNA. Recently, in H9 ESCs, an increase in the translation rate and differential methylation was observed in the FMR1 KO background, similar to the Shef4 hESCs, and these differences were predominant in the undifferentiated cells (Ninochka D’Souza *et al*., 2025). The intriguing differences in the methylation of rRNAs observed from the RMS data in the WT and the FMR1 KO ESC cells prompted us to ask if any inferences can be made from the structures of the ribosomes, including changes in their composition and conformations. Further, we asked if the changes in the 2’-O-methylation as observed in the RMS can be mapped between the two conditions. We employed cryo-electron microscopy (cryo-EM) of the ribosomes isolated from both the WT and the FMR1 KO ESCs and determined the structures from independent experiments. The cryo-EM analysis revealed that the ribosomes from the FMR1 KO background have a significant sub-population of dormant ribosomes marked by the presence of SERPINE mRNA binding protein (SERBP1), eukaryotic elongation factor (eEF2) or Ly-1 antibody reactive clone (LYAR) proteins along with translating ribosomes. Further, the methylation patterns of the WT and the FMR1 KO ribosomes were analysed in the 28S rRNA of these ribosomes but no large change in the ribosome structures was observed indicating that the differences in methylation have other distinct roles that might include local changes in interactions, but with the current resolution of the maps, it is difficult to expand on this particular feature.

## Results

### Structure determination of the wild type and the FMR1 KO ribosomes from hESCs

Embryonic stem cells typically have reduced translation rates and upon differentiation, the rate of translation increases (Saba *et al*., 2021). Global translational rates in cells were measured using fluorescent non-canonical amino-acid tagging (FUNCAT). As described previously for Shef4 hESCs, this assay revealed that the lack of FMRP due to FMR1 knockout resulted in an increase in translation in H9 ESCs **(Supplementary Figure 1)** (Ninochka D’Souza et al., 2025).

Ribosomes from both the WT H9 human embryonic stem cells (hESCs) and FMR1 KO hESCs were isolated after cycloheximide treatment and sucrose density gradient fractionation, followed by polysome profiling **(Figure 1a and 1b).** Initially, both the monosomes and the pooled polysome fractions were collected and observed by cryo-EM. However, very few molecules in the polysome fraction were observed and hence only the 80S fractions were further analyzed. The isolated 80S monosome fractions were pooled, centrifuged and the resuspended ribosomes were used for cryo-EM grid preparation and analysis. The structures of ribosomes from both the WT and the FMR1 KO cells were determined at an average overall resolution of ∼2.9 Å respectively after the initial 3D refinement **(Figure 1c and 1e).** The cryo-EM maps reveal well-resolved regions for much of the ribosomes, with only a few peripheral regions limited in resolution due to their inherent intrinsic flexibility (**Supplementary Figures 2 & 3**). The general architecture of ribosomes from hESCs is very similar to ribosomes previously purified from different mammalian cell lines, for example, HeLa cells (PDB 6qzp) **(Figure 1d and 1f).** The core ribosomal proteins typically found in other cell lines were observed but few polypeptides showed less resolved density. Some examples include the polypeptides S12, S27A, and uL1, whose density could be observed in the unsharpened maps but were smeared/fragmented after B-factor sharpening and only moderately resolved density was observed in the maps derived from deepEMhancer. Thus, these polypeptides are not included in the final deposited model, though they are present as part of the complex (**Table 1**). In the well-resolved part of the core regions, ions and water molecules could also be modelled with higher confidence. In addition, both base and ribose modifications were also modelled wherever the density was clear.

**Figure 1.**
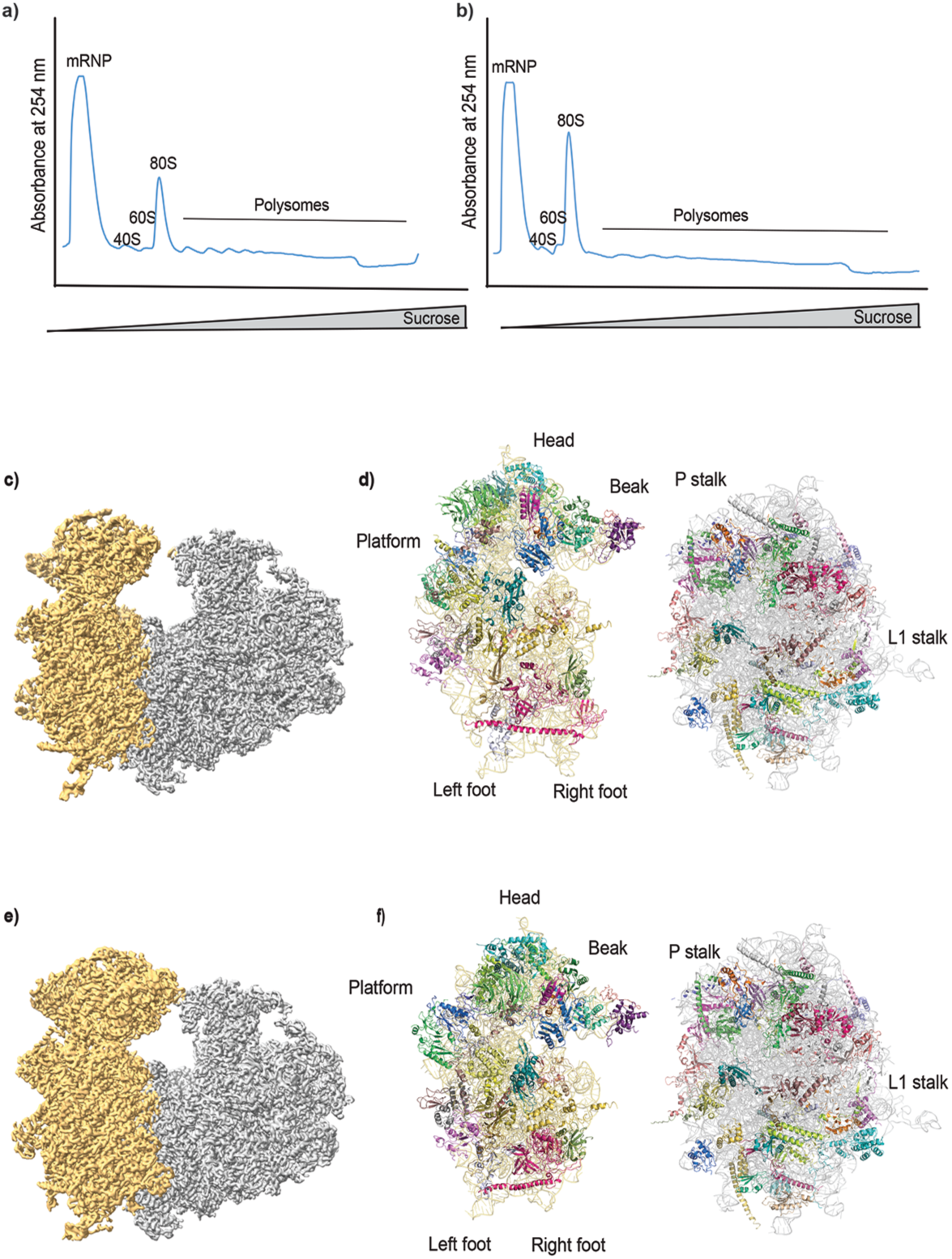
Cryo-EM maps of the WT and the FMR1 KO ribosomes from human embryonic stem cells (hESCs) **a) and b)** Polysome profile of the WT and the FMR1 KO hESCs respectively. The concentration of sucrose in the gradient ranges from 15-45 % and various fractions corresponding to mRNP, 40S, 60S, 80S (monosomes) and polysomes are labelled. The X-axis shows the increasing sucrose concentration and the Y-axis shows the absorbance at 254 nm. **c) and e)** Cryo-EM maps of the WT and the FMR1 KO ribosomes respectively. The deepEMhancer maps are shown at a threshold of 0.2 displayed in ChimeraX with a hide dust value of 30. The regions of the map corresponding to the large subunit are colored in gray and the small subunit in gold. **d) and f)** The models of WT and KO 40S and 60S subunits. The 18S rRNA is colored in gold, and the 28S rRNA is colored in gray and the different protein chains are shown in cartoon representation in both WT and KO ribosomes. Protein P0 (uL10) and L10A (uL1) have not been included in the model shown here as P0 is not found in most classes and L10A is highly dynamic with very poor density. Polypeptides S12 and S27a are included in the figure but not present in the final deposited model as they are less resolved.

**Table 1:** CryoEM Data collection, processing and Model statistics.

| CryoEM Data collection and processing | H9 ESC, WT Dataset 1 | H9 ESC, FMR1 KO, Dataset 1 SERBP1+eEF2 | H9 ESC, FMR1 KO, Dataset 1 with LYAR | H9 ESC, WT Dataset 2 | H9 ESC, FMR1 KO, Dataset 2 SERBP1+eEF2 | H9 ESC, FMR1 KO, Dataset 2 with tRNA |
| --- | --- | --- | --- | --- | --- | --- |
| EMDB ID | EMD-69344 | EMD-69362 | EMD-69345 | EMD-69346 | EMD-69347 | EMD-69348 |
| PDB ID | 23XA | 23YF | 23XB | 23XC | 23XD | 23XE |
| Magnification (nominal) | 75,000X | 75,000X | 75,000X | 75,000X | 75,000X | 75,000X |
| Voltage (kV) | 300 | 300 | 300 | 300 | 300 | 300 |
| Electron exposure (e<sup>-</sup>/Å<sup>2</sup>) | 27.5 | 25 | 25 | 25.4 | 25.6 | 25.6 |
| Defocus range (μm) | 1.5-3 | 1.5-3 | 1.5-3 | 1.5-3 | 1.5-3 | 1.5-3 |
| Pixel size (Å) | 1.07 | 1.07 | 1.07 | 1.07 | 1.07 | 1.07 |
| Symmetry imposed | C1 | C1 | C1 | C1 | C1 | C1 |
| Initial particle images (no.)<sup>a</sup> | 188015 | 135829 | 268650 | 190936 | 122650 | 122650 |
| Final particle images (no.)<sup>a</sup> | 149014 | 34045 | 220000 | 79956 | 67718 | 18877 |
| Map resolution FSC @0.143 (Å) | 2.9 | 2.9 | 2.5 | 2.9 | 2.9 | 3.1 |
| Map resolution range (Å)<sup>b</sup> | 2.5→5 | 2.5→5 | 2.2→6.5 | 2.5→5 | 2.5→5 | 2.5→5 |
| Map vs model FSC @ 0.5 (Å) | 3.1 | 3.16 | 2.8 | 3.0 | 3.0 | 3.2 |
| **Model Refinement** |  |  |  |  |  |  |
| Initial model used | H9, ESC, WT, Data set 2 | 6QZP/8acd/8qoi | 6QZP/8acd/8qoi | 6QZP/8acd/8qoi | H9, ESC, FMR1 KO, data set 1 | H9, ESC, FMR1 KO, data set 1 |
| Map sharpening *B* factor (Å<sup>2</sup>)<sup>c</sup> | -125.5 | -96.6 | -92 | -112.9 | -116.4 | -117 |
| **Model composition (# of atoms)<sup>d</sup>** |  |  |  |  |  |  |
| Overall | 207577 | 217607 | 208198 | 207632 | 217379 | 210755 |
| Protein | 90017 | 98909 | 90517 | 90017 | 98643 | 89752 |
| RNA | 117140 | 118256 | 117160 | 117140 | 118256 | 120609 |
| Ligands (ions) | 234 | 220 | 262 | 275 | 262 | 265 |
| Water | 186 | 222 | 259 | 200 | 218 | 129 |
| ***B* factors (Å<sup>2</sup>)** |  |  |  |  |  |  |
| Overall | 162.4 | 128.6 | 126.3 | 129.1 | 123.7 | 121.1 |
| Protein | 169.3 | 135.1 | 132.1 | 132.8 | 128.9 | 122.7 |
| RNA | 157.4 | 123.3 | 122 | 126.3 | 119.7 | 120.1 |
| Ligands | 110.1 | 79.2 | 91.8 | 92.9 | 85.9 | 85.4 |
| Water | 92.5 | 63.6 | 70.8 | 76.1 | 69.3 | 69.3 |
| **R.m.s. deviations** |  |  |  |  |  |  |
| Bond lengths (Å) | 0.008 | 0.008 | 0.008 | 0.008 | 0.008 | 0.008 |
| Bond angles (°) | 1.2 | 1.2 | 1.2 | 1.2 | 1.2 | 1.2 |
| **Validation** |  |  |  |  |  |  |
| MolProbity score | 1.2 | 1.3 | 1.2 | 1.2 | 1.2 | 1.2 |
| Clashscore | 0.6 | 0.9 | 0.6 | 0.6 | 0.8 | 0.5 |
| Poor rotamers (%) | 0.7 | 0.8 | 0.6 | 0.7 | 0.7 | 0.3 |
| **Ramachandran plot** |  |  |  |  |  |  |
| Favored (%) | 92.3 | 92.0 | 92.4 | 92.3 | 91.9 | 91.6 |
| Allowed (%) | 7.1 | 7.4 | 6.8 | 7.0 | 7.4 | 7.7 |
| Disallowed (%) | 0.6 | 0.6 | 0.8 | 0.7 | 0.7 | 0.7 |

<sup>a</sup> – the initial and final numbers are after classification. The workflow in supplementary figures has complete details on the number of images and particles.
<sup>b</sup> – the map resolution range is based on the local resolution
<sup>c</sup> – the values of map sharpening are from RELION post-process, for model building other maps with different B-factor sharpening or with deepEMhancer or LocSpiral were used.
<sup>d</sup> – the number of chains modelled (proteins and RNA) varies in different classes. The WT ribosomes from both the datasets and FMR1 KO dataset 1 with LYAR have 77 chains. LYAR though observed is not included in the final model due to very poor density. The FMR1 KO dataset 1\_eEF2+SERBP1 has 80 chains (77 chains as in WT and eEF2+SERBP1+uL10), KO\_dataset 2\_eEF2+SERBP1 has 81 chains modelled (77 chains as in WT and eEF2+SERBP1+uL10+RPL12) and KO\_dataset 2\_tRNA has 80 chains modelled (77 as in WT + 2xtRNA + mRNA).

### FMR1 KO cells have a subset of a dormant population of ribosomes

Following extensive 3D classification and multiple rounds of refinements, we found that in the WT cells, most of the ribosomes can be classified into different classes based on the global 40S movement but no additional proteins, tRNAs or mRNA bound to the ribosomes were observed, indicating that they are empty **(Figure 2a and Supplementary Figure 2)**. Surprisingly, in the ribosomes isolated from the FMR1 KO cells, we observed that there were at least two populations that had extra densities near the mRNA tunnel and RNA binding site and were proteinaceous in nature, along with a population with no factors bound **(Figure 2b and Supplementary Figure 3)**. To find the identity of these extra densities, an initial poly-alanine model was built (and with the aid of deepEMhancer and locSpiral maps) (Sanchez-Garcia *et al*., 2021; Kaur *et al*., 2021) and was identified as the eukaryotic elongation factor (eEF2) **(Figure 2c).** Based on the previous observations (Leesch *et al*., 2023; Wells *et al*., 2020; Brown *et al*., 2018), we identified the second factor as the mammalian SERPINE mRNA binding protein (SERBP1), which helps sequester eEF2. Both these factors inhibit mRNA and tRNA binding to the ribosomes and keep them “dormant” (Leesch et al., 2023; Wells et al., 2020). SERBP1 binds inside the mRNA tunnel whereas eEF2 binds at the interface near the A-site of the ribosome and interacts with the P-stalk. SERBP1 is a large protein (408 amino acids) with a high compositional bias and a highly disordered structure but in the map, only a small stretch is visible in the mRNA tunnel interacting with domain IV of eEF2 **(Figure 2d).** In the same population, we also observed the binding of eukaryotic translation initiation factor 5A (eIF5a) near the E-site. The presence of these factors prevents the binding of the tRNA and mRNA, thereby promoting the dormant state of the ribosomes.

**Figure 2.**
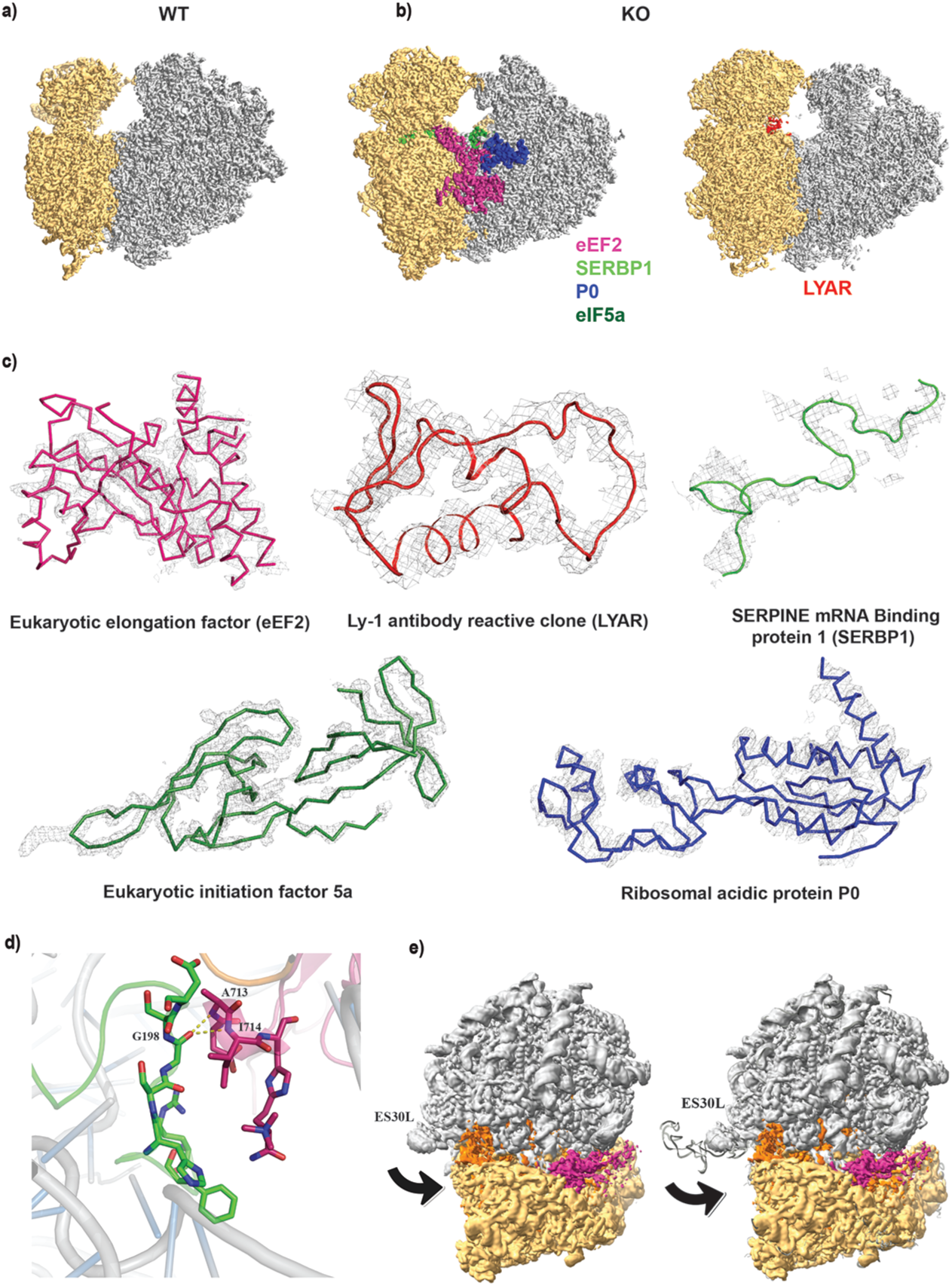
Distinct populations of ribosomes in the hESC WT and FMR1 KO cell lines in dataset 1. **a)** The 3D classification of WT hESCs revealed a single large population of ribosomes with no factors bound. The large subunit is colored in gray and the small subunit in gold. The deepEMenhancer map is shown at a threshold of 0.1 with a hide dust feature at 30. **b)** In the FMR1 KO cells, ribosome populations with different factors bound could be separated by 3D classification. One population had eukaryotic elongation factor (eEF2, magenta), eukaryotic initiation factor 5a (eIF5a, forest green), SERPINE mRNA binding protein (SERBP1, green) and acidic protein P0 (dark blue) along with other core ribosomal proteins. The deepEMhancer map has been contoured at 0.2 threshold with a hide dust value of 10. To individually color the additional subunits, a color zone radius of 10 in Chimera was used. A second distinct and abundant population had the cell growth-regulating nucleolar protein (LYAR, colored in red) bound near the interface of the large and the small subunit. The locSpiral map at a threshold of 5.75 with hide dust value of 5 and the color zone radius of 12 was used to show the LYAR density. **c)** To identify the factors bound in FMR1 KO ribosomes, poly-alanine models were built in the ordered regions, and this structural fold was used as input to the Dali server to identify the protein. The fit for the poly-alanine polypeptides or the final structures of the factors identified are shown in the respective cryo-EM maps. The first factor is the eukaryotic elongation factor (eEF2) where the density is carved out for the stretch between 93-354 residues showing a good fit. Few loop regions are flexible and the density for these regions is not very clear. The second factor is LYAR, for which the fit of the poly-alanine model used for identification is shown. SERBP1 was identified from the literature after identification of eEF2 and therefore, the part of the protein that is visible in the cryo-EM map has been modelled, and it fits in the mRNA tunnel. An initial poly-alanine polypeptide was built for both the Eukaryotic initiation factor 5a (eIF5a, density for modified lysine is clearly visible) and the ribosomal acidic protein P0 and fit of the full models is shown here. Densities for all the factors are carved at 2 Å from the atoms and ribbon representation is used for eEF2, eIF5a and P0, whereas, cartoon representation is used for LYAR and SERBP1. In the final deposited models, LYAR and eIF5a have not been included. **d)** SERBP1 binds in the mRNA tunnel and stabilizes the eukaryotic elongation factor (eEF2) near the A-site of the ribosome. The zoomed-in image shows the interface between SERBP1 and eEF2. SERBP1 is shown in green and eEF2 in magenta. The interacting residues are shown in stick representation while the neighboring regions are depicted in cartoon representation. SERBP1 contacts eEF2 near its domain IV, and potential interactions are mediated by the backbone of residue 198 of SERBP1 and 713-714 of eEF2. **e)** The movement of ES30L in different populations of the ribosomes is shown. The unsharpened final refined map, overlaid with a difference map has been used to demonstrate the movement. The extended part of the model, which is visible as part of ES30L, is usually flexible. The orange and pink densities are the difference densities present in dormant class. The pink color depicts the eEF2 and the orange highlights the change in the ES30L conformation (marked by black arrow) when all the factors (eEF2+eIF5a+SERBP1+P0) are present. In the left panel, the model has been overlaid to show the protruding ES30L. The map with exposed ES30L is at the threshold of 0.022, the dormant class is at a threshold of 0.055 and the difference volume is at a threshold of 0.054.

The population of ribosomes with eEF2, SERBP1 and eIF5A exhibits a conformational change of the ES30L expansion segment, which is a 62-nucleotide-long (3975-4036) extension of helix 78 of 28S rRNA and is part of the L1 stalk in humans. ES30L is highly dynamic and lies in a solvent-accessible region of the ribosome, consequently, the cryo-EM maps in this region typically exhibit poor density (Natchiar *et al*., 2017; Khatter *et al*., 2015; Anger *et al*., 2013). We observed that the ES30L segment is solvent-exposed, and flexible in all the classes, consistent with previous structures, but in the dormant class containing all three factors, it is oriented toward the E-site of the ribosome **(Figure 2e).**

In a substantially large population of this dataset, we observed another density bound near the A/P-site of the ribosome, and after de novo poly-alanine model building and identification, it was identified as the cell growth-regulating nucleolar protein or Ly-1 antibody reactive clone (LYAR) **(Figure 2c).** LYAR has been observed previously in sub-populations of ribosomes isolated along with Nsp1-80S complexes (Thoms *et al*., 2020), with endothelial differentiation-related factor 1 (EDF1) associated ribosomes (Sinha *et al*., 2020) and, more recently, in the ribosomes from MCF-7 cell line, where it is found in the decoding center (Trendel *et al*., 2022). It is clear from the location of LYAR that it can prevent translation but its relevance in the FMR1 KO background is unclear at the moment. Higher translation rates are observed in the FMR1 KO ESCs when compared to the WT ESCs, but it is interesting that these cells also have ribosome populations that are dormant **(Supplementary Figure 1).** The isolation of the ribosomes from both the WT and the FMR1 KO cells was performed simultaneously using the same protocol, but there is a possibility that some of these factors observed in the FMR1 KO ribosomes were lost during the enrichment process.

Therefore, to validate our results obtained from the first dataset, a separate batch of cells was grown and subjected to ribosome purification followed by cryo-EM analysis as described above, which we refer to as dataset 2 **(Figure 3).** In this replicate, we again observed largely empty ribosomes from WT H9 ESCs similar to the first dataset **(Figure 3a and Supplementary Figure 4).** When the FMR1 KO ribosomes were subjected to 3D classification, two major classes (class 2 and class 4) were identified: one with clear density for eEF2 and the other with residual density in the mRNA/tRNA-binding site **(Figure 3b and Supplementary Figure 5).** These two major classes were further subjected to 3D classification, and several sub-populations could be separated with eEF2 and SERBP1 (class 4_1 and class 4_3) or with tRNA bound (class 4_2 and class 4_4). These populations of the ribosomes with tRNAs reveal that they are in the A and P sites and in the P and E sites respectively **(Figure 3c and 3d).** Class 4_1 with eEF2 and SERBP1 and class 4_4 with two tRNAs were used for further analysis as the density in the other two classes was relatively poor.

**Figure 3.**
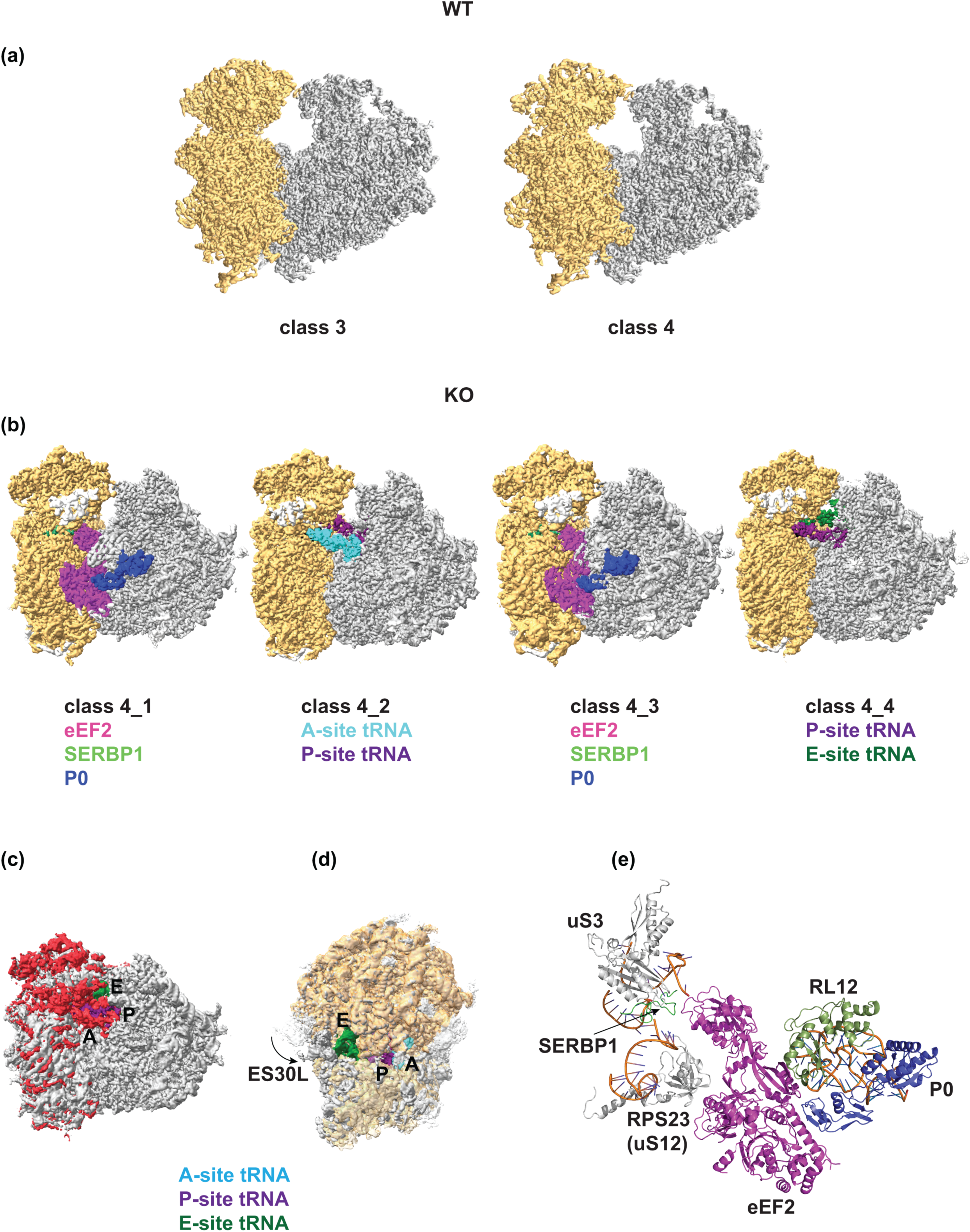
Distinct populations of ribosomes in the hESC WT and FMR1 KO cell lines in dataset 2. **a)** In dataset 2 of the WT hESCs, 2 populations of ribosomes with difference in the global movement of 40S were separated after 3D classification. The large subunit is colored in gray and the small subunit in gold. For both classes, deepEMhancer maps at a threshold of 0.2 with hide dust value 30 was used. **b)** In the FMR1 KO cells, populations of both dormant and tRNA containing ribosomes were obtained. In the two dormant populations obtained, eukaryotic elongation factor (eEF2, deep pink), SERPINE mRNA binding protein (SERBP1, green) and acidic protein P0 (dark blue) were observed but no eIF5a (unlike dataset 1). Both these classes varied in the 40S global movement. The other two classes in this dataset had tRNAs. One of them had the A-site tRNA (cyan), the P-site tRNA (purple) and the mRNA (black), whereas, the other population had the P-site tRNA (purple), the E-site tRNA (green) and the mRNA (black). The deepEMhancer maps at thresholds varying between 0.08 and 0.2 have been used for depiction, where the subunits are individually colored using the color zone option in ChimeraX with the zone radii of 6. **c)** The difference in class 4_2 and class 4_4 is shown here with calculation of a difference map (with unsharpened map without normalization in ChimeraX). The map in red is the difference map (class 4_4-class 4_2,) and overlaid on class 4_4 map with the P and E-site tRNAs colored. The position of the A-site tRNA is marked. **d)** An overlay of the class 4_2 (shades of yellow for the 60S and 40S subunits) and class4_4 (gray), the tRNAs in both the maps are colored as in panel b. The overlay shows the 40S movement, which is the red difference density in panel c. Note that the tRNA molecules could be in hybrid A/P and P/E sites in class 4_2, currently labelled as A and P sites. The movement of ES30L is marked with an arrow and this is unmodelled. **e)** The interaction of P0 and SERBP1 with regions of the ribosomes. The view and coloring are as in panel b. The C-terminus of P0 interacts with domain I of eEF2 and the N-terminus interacts with 28S rRNA and RPL12 (uL11). Domain IV of eEF2 interacts with SERBP1, which is surrounded by 18S rRNA and 40S subunits uS3 and RPS23 (uS12). For clarity, only few proteins and segments of RNA near SERBP1 and eEF2 are shown.

Interestingly, in both the populations with eEF2 and SERBP1 (class 4_1 and class 4_3), no eIF5a was observed but the proportion of the ribosomes with eEF2 and SERBP1 (>50%) was much larger than in dataset 1 (∼5%). Notably, in the population with eEF2/SERBP1, the conformational change of ES30L was not observed perhaps due to the absence of eIF5A **(Figure 2e).** In this map, another extra density was observed and it was identified as acidic protein P0, which along with other acidic proteins is known to form the P-stalk of the ribosome and helps in translational elongation **(Figure 3b and 3e)**. Later, the density for P0 was also identified in dataset 1, which was most likely missed due to its lower occupancy. In the current workflow, the analysis is limited to the purified ribosomes and not the total ribosomes in cells, and the populations observed are limited by the experimental procedure, for instance, LYAR or eIF5A, and tRNA bound populations were observed in one of the data sets and not in both. Nevertheless, the two independent data sets of ribosomes isolated from FMR1 KO cells reveal substantial dormant-like subpopulations of ribosomes, while the ribosomes from WT preparations largely contain empty ribosomes with no factors bound.

### Localization of 2’-O-methylations in the WT and the FMR1 KO hESCs

One of the intriguing functions of FMRP is its interaction with snoRNAs and its nuclear localization, which results in differential 2’-O-methylation of both 28S and 18S rRNAs as evident from RMS analysis (Ninochka D’Souza *et al*., 2025; D’Souza *et al*., 2018). The RMS data reveal hypermethylation or hypomethylation of certain residues and not a complete loss, which makes structural interpretation a difficult proposition and requires maps that are well-ordered and at a sufficiently high resolution. We used multiple maps with different levels of sharpening to identify the differentially methylated sites and also the difference maps calculated by servalcat (**Supplementary Figure 6**) (Yamashita *et al*., 2021). In validating the methylation sites, we also inspected the density of the neighbouring residues for an unbiased assessment and also verified all the potential methylations and not just the ones highlighted by the RMS data. Further, we checked if the maps at the regions of interest were of sufficient quality by mapping the local resolution values along with the refined B-factor values (**Figure 4a and 4b and Supplementary Figures 2, 3, 4, 5**). Most of the sites identified from the RMS studies were found to be methylated in 28S rRNA of all the classes, and one such example is 1522, 2’-O-methylguanosine (OMG), which is part of helix 32 **(Figure 4c).** Of the methylated residues analysed in the 28S rRNA, residue 4618 which forms part of helix 95 could be mapped as differentially methylated between the WT and the FMR1 KO ribosomes with some confidence **(Figure 4d).** When the density for the methyl group was analysed at different contour levels for residue 4618, it was better resolved (or showed higher occupancy) in the FMR1 KO ribosomes, whereas the methylation showed lower occupancy in the WT ribosomes **(Supplementary Figure 7)**. The mapped methylations on the 28S rRNA of the WT and the FMR1 KO ribosomes are shown in Figure 4e. The sub-populations of each dataset had varying numbers of particles and resolution, but still the methylation patterns were consistent **(Figure 4 and Table 1 and Supplementary Table 1).** With the current data, we could not identify any residue that had undergone hypomethylation; that is, no sites showed the opposite trend of methylation between WT and FMR1 KO cells (i.e., methylated in WT and unmethylated in KO with high confidence; **Supplementary Table 1**). This is not unexpected as FMRP has been reported to interact with specific snoRNAs and may thereby influence rRNA methylation patterns (D’Souza *et al*., 2018; Ninochka D’Souza *et al*., 2025). On the other hand, the differentially methylated residues of the 18S rRNA from the RMS are in the highly flexible regions and thus, methylation differences in the 40S subunit were not analysed because they could not be mapped confidently, but the well-studied di-methylated adenines (m^6^_2_A 1850 and m^6^_2_A 1851) could be modelled.

**Figure 4.**
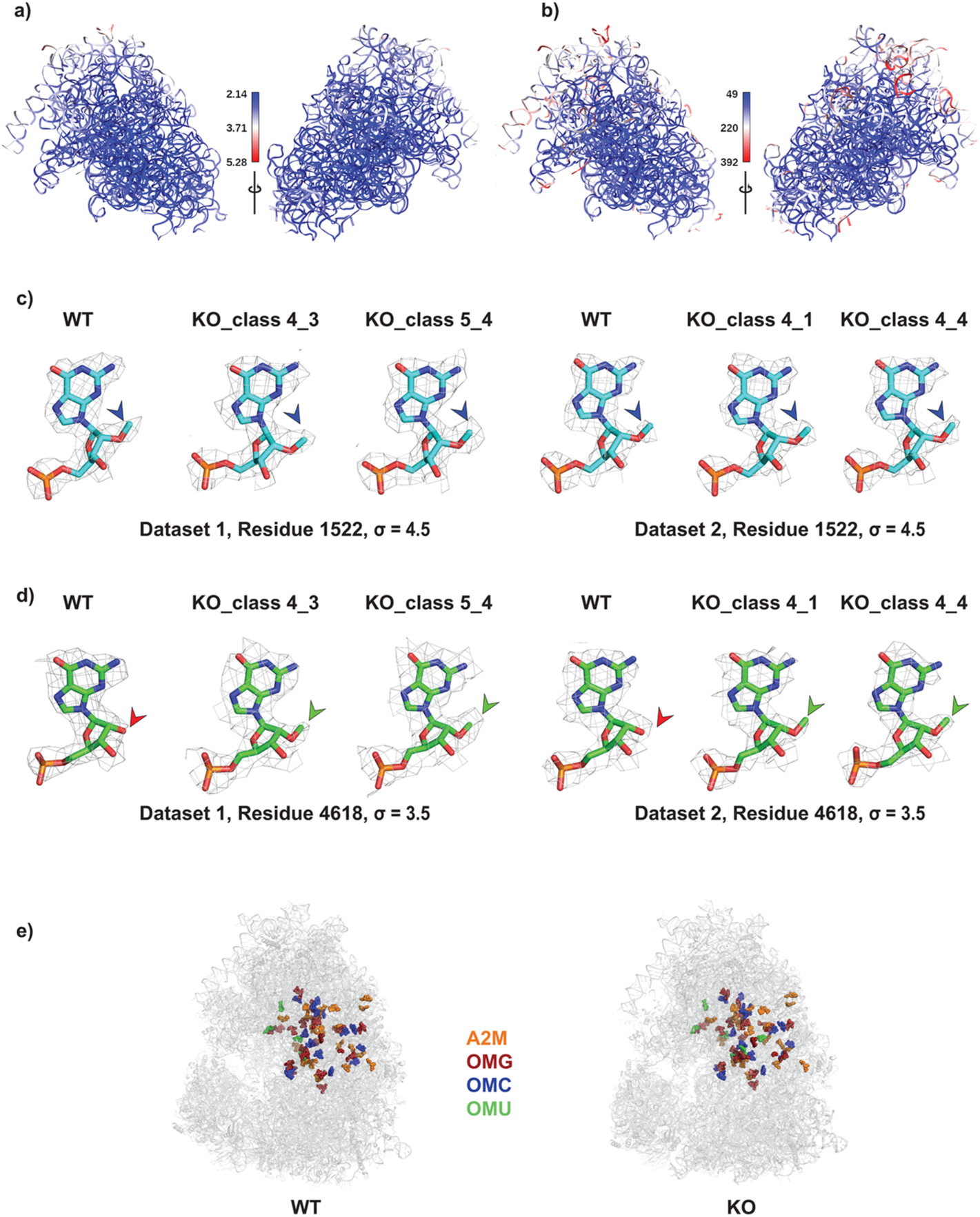
Mapping 2’-O-methylation in the hESC WT and the FMR1 KO ribosomes. **a and b**) Two views of 28S rRNA model (FMR1 KO, data set 1, population with LYAR, class 5_4) coloured according to the map values estimated with LocalRes, and with B-factor. The range of the local resolution (Å), and the B-factor (in Å^2^) are shown. Similar plots for other populations in the two data sets are shown in supplementary figures 2-5. The map of 28S rRNA coloured according to the local resolution with the methylated residue is shown in supplementary figure 6. **c)** Residue 1522 of 28S rRNA showing 2’-O-methylation in all the populations. The residue 1522 has a methyl group at 2’-O of the ribose (blue arrow) in both the WT and the FMR1 KO ribosomes, and is present in all the different populations of both the datasets. The cryo-EM density is shown in gray, mesh at σ=4.5 and carved around 1.75 Å from the atoms. **d)** Residue 4618 that is identified to be differentially methylated by RMS is shown from both the WT and the FMR1 KO ribosome datasets to depict the absence of methylation in WT (red arrow) and presence of a clear density for the methyl group in the FMR1 KO ribosomes (green arrow). The cryo-EM density is carved 1.75 Å around the atoms with the σ=3.5. **e)** Location of all the 2’-O-methylations that have been modelled in the WT and the FMR1 KO ribosomes respectively. A2M (orange spheres) marks the 2’-O-methylation on adenine residues, OMG (firebrick spheres) marks the 2’-O-methylation on guanine, OMC (blue spheres) marks the 2’-O-methylation on cytosine residues and OMU (green spheres) marks the 2’-O-methylation on uracil residues on WT and FMR1 KO ribosomes respectively.

For an independent quantitative measure, MapQ was used to analyse the resolvability of the methyl groups in the maps (Pintilie *et al*., 2020). For this analysis, methyl groups were added at the 2’-O of the ribose in residue 4618 in both WT datasets and in residues 1323 and 3867 in the FMR1 KO dataset 2 with tRNA. Visual inspection of these residues in the maps revealed no clear density for the methyl groups. While the MapQ values for residues 1323 and 3867 were lower in the FMR1 KO dataset 2 with tRNA, the value for residue 4618 was surprisingly similar in the WT and FMR1 KO ribosomes (**Supplementary Table 2**). Indeed, when the contour level is reduced then residual density for residue 4618 in the WT ribosomes could be observed (**Supplementary Figure 7**). This highlights the limitation in identifying small changes in methylation patterns from the cryo-EM maps, and thus analysis of hyper/hypo methylation requires other quantitative methods.

## Discussion

Due to their lack of a defined molecular identity, ESCs have been studied as model systems. They are known to repress cell type-specific genes, while expressing genes that maintain their pluripotency. ESCs respond to the environmental cues to change their cellular identity and in this process, a quick cellular response increases the rate of protein translation while the undifferentiated ESCs have low translation rates (Gabut *et al*., 2020; Sampath *et al*., 2008). When the H9 ESCs are differentiated into neural stem cells by inhibiting the SMAD signalling pathway, the translation rate increases and differential methylation is observed (Ninochka D’Souza *et al*., 2025). The role of FMRP as a translational regulator and in modulating the rate of protein synthesis inside a cell is well-studied (Thelen & Kye, 2020). The primary mechanism of translation regulation is via binding to ribosomes (E. Chen et al., 2014), but binding to C/D box snoRNAs is another means of regulation (D’Souza *et al*., 2018). Previous studies have shown that loss of FMRP not only results in an increase in translation **(Supplementary Figure 1)** but also changes in the methylation of rRNA (D’Souza *et al*., 2018; Ninochka D’Souza *et al*., 2025). Since knockouts of snoRNA are technically challenging in hESCs, FMR1 KO hESCs provide a good model system to understand the effect of change in methylation and translational regulation, which forms the basis of this study.

Structural analysis of the WT and FMR1 KO ribosomes from human embryonic stem cells (H9 hESCs) provides significant insights into their architecture and additional factors associated with core ribosomes **(Figure 1d and 1f**, **Figure 2a and 2b and Figure 3a and 3b).** As anticipated, the core structure of ribosomes is consistent with previously determined structures from other human cell lines, reinforcing the conserved nature of ribosome architecture across different human cells (Anger *et al*., 2013; Khatter *et al*., 2015; Du *et al*., 2024). While the ribosomes from the WT H9 ESCs were observed to be in an empty state without any bound factors in two different data sets, the ribosomes from FMR1 KO preparations revealed different populations. In the first data set, we observed two dormant populations of ribosomes in FMR1 KO H9 ESCs, which had either a combination of SERBP1, eEF2 and eIF5A or LYAR **(Figure 2b).**

LYAR is a zinc finger protein and is known to associate with nucleolin, and thought to be important for maintaining ESC identity (Li *et al*., 2009). In the context of ribosomes, LYAR has previously been observed in studies involving SARS-CoV2 infection, structural analysis of collided ribosomes and in understanding the ribosome lifecycle (Trendel *et al*., 2022; Sinha *et al*., 2020; Thoms *et al*., 2020), and we have observed LYAR to be present in some ribosomal preparations of HEK 293 cells (unpublished data). All these reports are from different cell lines. It is not clear to us in what physiological context LYAR binds to the ribosomes and it is even more intriguing as it is thought to bind to nucleolin and resides in the nucleus (Li *et al*., 2009). Nevertheless, in our map as well as in the published structures, LYAR occupies around the A-site and P-site, and likely prevents translation, thus making it an interesting ribosome associated factor that is less studied physiologically (**Figure 2b**).

Several cellular factors that bind to ribosomes and promote dormancy in eukaryotes are now well documented and these include IFRD2, CCDC124, Lso2 and SERBP1, and its yeast homologue, Stm1 (Smith *et al*., 2022). The physiological contexts of such dormancy vary from nutrient starvation, differentiation (A. Brown et al., 2018; Kišonaitė et al., 2022, Leesch et al., 2023; Van Dyke et al., 2006, 2013; Wells et al., 2020), and some viral proteins can also repress host translation (Thoms *et al*., 2020). Upon a stimulus and differentiation, ESCs undergo rapid change and the rate of translation increases to become a defined cell. Such an increase in translation is also observed in FMR1 KO ESCs, highlighting a potential role for translational regulation mediated by FMRP. Thus, a possible scenario is that FMRP perhaps acts as a brake either by directly binding to ribosomes or indirectly by binding to mRNAs (E. Chen et al., 2014; Richter & Zhao, 2021; Thelen & Kye, 2020). However, the increase in translation observed by FUNCAT in FMR1 KO cells (**Supplementary Figure 1**), and the dormant populations observed by cryo-EM analysis cannot be correlated with the current data. That is, the loss of FMRP resulting in increased translation prompting the cells to cycle some ribosomal populations to dormant state with conserved protein factors (**Figure 2, 3**). The absence of FMRP might be regarded as a stress to the cells, and the presence of dormant ribosome populations in the purified FMR1 KO ribosome preparations might be reflective of this. Further, the proportions of the subclasses observed in the datasets are not reflective of total ribosomes in the cells, and thus a direct mechanistic link cannot be established between the loss of FMRP and the observation of dormant ribosomes from the analysis described here.

In other eukaryotic cells, *in situ* studies of ribosomes show populations of dormant ribosomes and cellular states such as stress might promote such dormancy (Gemmer *et al*., 2023). Whether the absence of FMRP induces some stress that contributes to the dormant populations of ribosomes observed here remains to be determined. Mechanistically, how these proteins bind to assembled ribosomes, particularly long and disordered proteins such as SERBP1, remains an interesting question to be addressed.

The importance of ribose 2’-O-methylation in rRNA is now well documented in translation dynamics, development and cancer (Khoshnevis *et al*., 2022; Ayadi *et al*., 2019; Erales *et al*., 2017; Decatur & Fournier, 2002; Natchiar *et al*., 2017). The methylations of 2’-OH of the ribose moiety are known to be crucial for ribosome function and are observed in all ribosomes across the kingdoms (Ayadi *et al*., 2019; Decatur & Fournier, 2002; Natchiar *et al*., 2017). The mechanisms of how differences in methylation cause translation regulation are currently unclear. Studies in yeast have shown that reduction in these modifications affects the inherent dynamics of ribosomes and impacts the binding of translational regulators, thereby causing defects in protein synthesis (Khoshnevis *et al*., 2022).

As we determined the ribosome structures from the WT and FMR1 KO ESCs, we also analysed the methylation pattern in multiple populations of the ribosomes **(Supplementary Table 1).** In recent times, the achievable resolution of ribosome maps by cryo-EM has been steadily increasing with the culmination of <2 Å for human ribosomes (Holvec *et al*., 2024; Wiechert *et al*., 2025; Faille *et al*., 2023; Pellegrino *et al*., 2023). Such resolutions allow for modelling modifications in bases, sugar moieties, protein residues and modelling of ions and solvent molecules with greater confidence. The cryo-EM maps of ribosomes from hESCs described are of sufficient resolution to map the 2’-O-methylations on the ribose moiety in the well-ordered regions in both the WT and the FMR1 KO ESC ribosomes and pinpoint the differences **(Figure 4 and Supplementary Figures 2-5).** Unbiasedly all the residues of rRNA were cross-checked for methylation with the high-resolution structures (Faille *et al*., 2023; Pellegrino *et al*., 2023; Holvec *et al*., 2024; Wiechert *et al*., 2025). Although the maps for these structures are at very high-resolution, we observe at least two differences in the assignment of 2’-O-methylation **(Supplementary Figure 8).** These differences may be indicative of the cell-type differences as Expi293 and HeLa cells were used for ribosome preparations (Holvec *et al*., 2024; Faille *et al*., 2023; Wiechert *et al*., 2025). Residue 4618 showed a significant change in the RMS signal in the WT and the FMR1 KO cells, and therefore, was confidently modelled as differentially methylated in our structures, whereas, some other residues with a similar change in RMS are still ambiguous and have not been modelled (Ninochka D’Souza *et al*., 2025). These challenges also indicate the technical limitations and the lack of means to quantify these methylations structurally, and to correlate with RMS and thus alternative strategies such as mass spectrometry, for confirming these methylations are required.

The addition of a methyl group to the ribose of a nucleotide is extremely small, such minor changes in a large molecule like a ribosome can make a significant functional difference. It is known that modifications in mRNA and tRNA can regulate translation, for example tRNA-derived factors (tRFs) in stem cell function (Saba et al., 2021), and differential methylation in rRNA although less well studied, is gaining attention. Such an effect has been shown in HeLa S3 cells, where ribose methylation of C174 residue in 18S rRNA was shown to regulate the translation of specific mRNAs depending on the codons, although the residue is observed close to helix 8 of the small subunit further away from any catalytic site (Jansson *et al*., 2021). Thus, it is highly probable that the hypo and hyper methylations of ribose moiety would impact selective mRNAs and the dynamics of ribosomes like previously rather than having drastic structural effects.

In summary, these observations point towards the complex regulatory role of FMRP in translation and ribosome function. The presence of dormant subpopulations in FMR1 KO ESCs ribosome preparations underscores the necessity for a tightly controlled balance between translation activity and ribosome conservation, but the exact mechanistic role requires further studies. Understanding this balance could have broader implications for understanding the cellular economy of ribosomes and the pathological consequences of FMRP deficiency.

## Methods

### Embryonic stem cell culture

H9 ESCs and FMR1 KO ESCs were cultured on Matrigel (#3545277, BD Biosciences) coated plates containing mTeSR1 medium (#5850,StemCell Technologies) at 37°C in a 5% CO_2_ environment. Cells were further passaged with an enzyme cocktail containing 1 mg/ml of Collagenase type IV (#17104019, Invitrogen), 20% KOSR (#10828010,Gibco), 0.25% Trypsin and 1 mM CaCl_2_ dissolved in 1X PBS without CaCl_2_ or MgCl_2_ pH 7.2. For immunostaining experiments, H9 ESC colonies were plated on Matrigel-coated glass coverslips and cultured as mentioned above.

### Metabolic labelling and imaging

Embryonic stem cells (ESCs) were first incubated in methionine-free Dulbecco’s Modified Essential Medium (Thermo #21013024) for 30 minutes, followed by the addition of azidohomoalanine (AHA; 1μM, Thermo #C10102) in the same medium. After 30 minutes of incubation, the cells were fixed using 4% paraformaldehyde (PFA) for 10 minutes. Next, they were permeabilized with a PBS solution containing 0.3% Triton X-100 and then blocked with a buffer containing PBS, 0.1% Triton X-100, 2% BSA, and 4% FBS. Newly synthesized proteins were labelled with Alexa Fluor 555-alkyne [Alexa Fluor 555 5-carboxamido-(propargyl), bis (triethylammonium salt); Thermo #A20013] through a click chemistry reaction using the CLICK-iT cell reaction buffer kit (Thermo #C10269), allowing the fluorophore alkyne to react with the azide group of AHA. Cells were further immunostained for α-tubulin to identify cell structures. Finally, the coverslips were mounted using Mowiol® mounting medium (Sigma #81381).

The mounted coverslips were imaged using an Olympus FV3000 confocal laser scanning inverted microscope equipped with a 20X objective. To ensure proper sampling according to Nyquist’s criteria in the XY direction, the pinhole was set to 1 Airy Unit and the optical zoom to 2X. The objective was moved in the Z-direction with a step size of 1 μm, capturing approximately 8-10 Z-slices to collect light from planes above and below the focal plane. For FUNCAT imaging, cells were identified using the α-tubulin channel. Image analysis was performed using ImageJ software, where maximum intensity projections of the slices were used to quantify mean fluorescent intensities. Regions of interest (ROIs) were drawn around the cells based on the α-tubulin channel. The data are presented as box plots that show the quantification of FUNCAT fluorescent intensity normalized to the α-tubulin fluorescent intensity. The box plot extends from the 25th to the 75th percentile, with the median indicated by the middle line. Whiskers represent the full range of data from minimum to maximum values. Statistical significance was calculated using Unpaired Student’s t-test (2 tailed with equal variance).

### Polysome profiling

Human ESCs cells were lysed using the lysis buffer (20 mM Tris-HCl, pH 8.0, 100 mM KCl and 5 mM MgCl_2_, 1 mM DTT and 0.1 mg/mL CHX and 1% NP-40) and incubated at room temperature for 10 minutes on a shaker. The cell debris was separated by centrifuging the lysate at 21130*g for 20 minutes at 4 °C. The supernatant was then loaded onto preformed, pre-chilled 15-45% (w/v) linear sucrose gradients made in the gradient buffer (20 mM Tris-HCl, pH 8.0, 100 mM KCl, 5 mM MgCl_2_, 0.1 mg/mL CHX and 1 mM DTT), and ultra-centrifugation was performed at 39000 rpm for 90 minutes at 4 °C in Beckman Coulter’s SW40 rotor. After the centrifugation, the sample was fractionated into 12, 1 mL fractions with continuous UV absorbance measurement (A254) using 60% sucrose on a Teledyne ISCO polysome profiler. The fractions containing monosomes were pooled and ultracentrifuged at 100000*g for 1 hour at 4 °C to remove sucrose. The resulting ribosome pellet was resuspended in ribosome resuspension buffer (20 mM Tris-HCl, pH 8.0, 100 mM KCl and 5 mM MgCl_2_, 1 mM DTT) and used for grid preparation and cryo-EM analysis. For grid preparations, the samples were diluted to 200-250 nM final concentration.

### Cryo-EM Sample preparation and Data acquisition

Thin carbon film on mica was evaporated using an Edwards benchtop 150 carbon evaporator. The carbon was floated on Quantifoil R2/2 Au 300 mesh grids using a floating apparatus one day before cryo-EM sample preparation. On the day of freezing, Vitrobot Mark IV (ThermoFisher Scientific) was first set at 4 °C and 100% humidity. The grids were then glow-discharged using Quorum GloQube at 20 mA for 15 seconds and in the meantime, ethane was liquified. After glow discharge, the grids were mounted on the Vitrobot tweezers, 3 μL of freshly prepared ribosome sample was applied, held for 10 seconds and blotted for 3-4 seconds (blot force set to 0). The grids were plunged in liquid ethane after blotting and stored under liquid nitrogen conditions until imaging.

On the day of data collection, the grids were clipped into Titan Krios cartridge and loaded onto a 300kV Titan Krios G3i microscope (ThermoFisher Scientific). The data acquisition was performed using EPU software (ThermoFisher Scientific). Image shift and beam alignment were done before setting up data acquisition. The gain reference for the Falcon3 detector in integration mode was also acquired before each data collection. The datasets were collected in integration mode at a magnification of 75000X, corresponding to a pixel size of 1.07 Å, with a dose of 1-1.2 e^-^/Å^2^/frame and 20-25 movie frames were collected. Three images were collected from each hole and the data were acquired at a defocus range of -1.5 to -3.0 μm.

### Image processing

All the data processing steps were performed in RELION 3.1 (Scheres, 2012) except one of the FMR1 KO datasets which was processed with RELION 4.0 (Scheres, 2012; Zivanov *et al*., 2020, 2019; Kimanius *et al*., 2021). Multi-frame movies were motion-corrected and summed using Relion’s own MotionCor algorithm with the number of patches set to 5x5 in the X and Y-direction respectively, followed by contrast transfer function estimation using CTFFIND4 using FFT box size set to 1024 pixels (Rohou & Grigorieff, 2015). Subsequently, the micrographs were selected based on defocus values and CTF resolution estimation, or occasionally they were manually curated. Particles were picked using reference-free methods such as the Laplacian-of-Gaussian blob-detection method using a diameter range of 250-375 Å with a default threshold (stddev) of 1.2, followed by particle extraction with a box size of 384 pixels (360 pixels for WT, dataset 1). Post-particle extraction, 2D class averaging was performed iteratively to get high-resolution 2D class averages with a mask diameter of 340 Å. An initial 3D reconstruction was generated from the selected classes and further 3D refinement was done with C1 symmetry. This was followed by CTF refinement and Bayesian polishing to correct for beam tilt, trefoil, per particle defocus and astigmatism. Another round of 3D refinement was performed followed by post-processing with the half-maps to obtain sharpened maps. Extensive 3D classification was performed for all the data sets to identify sub populations with factors bound (described in the workflow) with different parameters including default settings, with mask and no alignment and the classes were inspected in chimera and the best classes were refined again. All the resolution estimation was done using gold-standard Fourier shell correlation at 0.143 after masking using the postprocess option in RELION. Local resolution was calculated with RELION.

### Model building and refinement

For model building, multiple B-factor sharpened maps were used in tandem with auto B-factor sharpened maps along with the unsharpened maps. In addition, deepEMhancer and locspiral were used to post-process the maps, thereby, improving their interpretability (Sanchez-Garcia *et al*., 2021; Kaur *et al*., 2021). Firstly, 6zmi and 6qzp human ribosome structures (and later 8acd and 8qoi) from the Protein Data Bank were docked into the map using the Chimera fit-in-map feature and the model was then saved relative to the map (Pettersen *et al*., 2021, 2004). The saved model was then opened in Coot superimposed on the map and local alignments were done to fit all the chains correctly (Emsley *et al*., 2010). The chains for which density was not clear or completely absent, were removed. After accounting for the core ribosomal subunits, other unmodelled densities were identified by building a poly-alanine chain and based on the fold, the identity of the chain was determined using the Dali server (Holm *et al*., 2023). Water molecules and ions were modelled based on their density and chemistry only in the well resolved regions. The cif files of the modified bases were generated using Jligand (Lebedev *et al*., 2012), and the modifications were added wherever the density was clear. For refinements of the model, appropriate CIF files were added as inputs along with the map and the model. The bundles containing different regions of the ribosomes were refined separately, combined after refinement, and geometry minimization was done before finally refining the combined model using Phenix and Refmac/Servalcat (Afonine *et al*., 2018; Adams *et al*., 2012; Murshudov *et al*., 2011; Yamashita *et al*., 2021). The final refinement was carried out with the half maps as input using Refmac/Servalcat (Yamashita et al., 2021). The output models were checked for bond angle, bond length deviations, Ramachandran outliers, rotamer outliers, clash score, etc using Phenix molprobity analysis (Williams *et al*., 2018; Chen *et al*., 2010). Although the polypeptides eIF5a and LYAR were identified in the subclasses of the FMR1 KO dataset 1, they were not included in the final model because of poor density. To map the local resolution onto the models, map values from local resolution mrc file estimated from RELION were used, and the attribute values were written to the B-factor column in the PDB file (only for the 28S rRNA as shown here), and used to colour the model in the respective local resolution range. For colouring according to the B-factor, the final refined model was used. These figures were made with ChimeraX (Pettersen *et al*., 2021).

## Acknowledgements

We acknowledge the National Cryo-EM facility, Bangalore, for data collection, which was supported by the Department of Biotechnology, DBT/PR12422/MED/31/287/2014, and the computing facility in the Bangalore Life Science Cluster in particular Mr Chakrapani. TK acknowledges the graduate fellowship from TIFR/NCBS. K.R.V. acknowledges the support of the Department of Atomic Energy, Government of India, Project Identification No. RTI 4006. KRV is part of the EMBO Global Investigator Network.

## Data Availability

The cryo-EM maps and the coordinates will be deposited in the EMDB and PDB respectively with the following accession codes.

H9 WT ESC, dataset 1– EMD-69344 and PDB 23XA

H9 WT ESC, dataset 2– EMD-69346 and PDB 23XC

H9 FMR1 KO, dataset 1, eEF2, SERBP1 - EMD-69362 and PDB 23YF

H9 FMR1 KO, dataset 1, LYAR - EMD-69345 and PDB 23XB

H9 FMR1 KO, dataset 2, eEF2, SERBP1 - EMD-69347 and PDB 23XD

H9 FMR1 KO, dataset 2, translating ribosome - EMD-69348 and PDB 23XE

