## Supplementary Information for "Analysis of ribosomes from WT and FMR1 KO human embryonic stem cells"

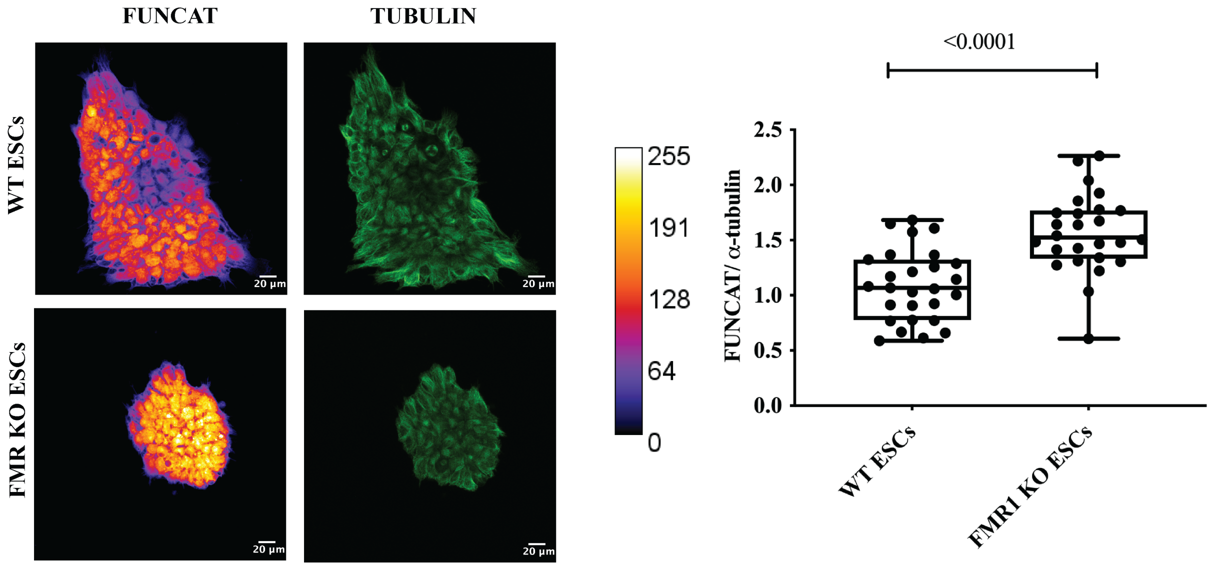


**Supplementary Figure 1. FUNCAT analysis of the WT and the FMR1 KO H9 hESCs**

**a)** Representative images for FUNCAT and α-Tubulin fluorescent intensities in the WT and the FMR1 KO ESCs. (Scale bar— 20 µm) **b)** Box plot representing the quantification of the FUNCAT intensity normalized to α-Tubulin intensity for each colony of the WT ESC and the FMR1 KO ESC. The box extends from the 25th to the 75th percentile with the middle line representing the median of the dataset. Whiskers range from minimum to maximum data points. Unpaired t-test. n=26-29 colonies from 3 independent experiments.

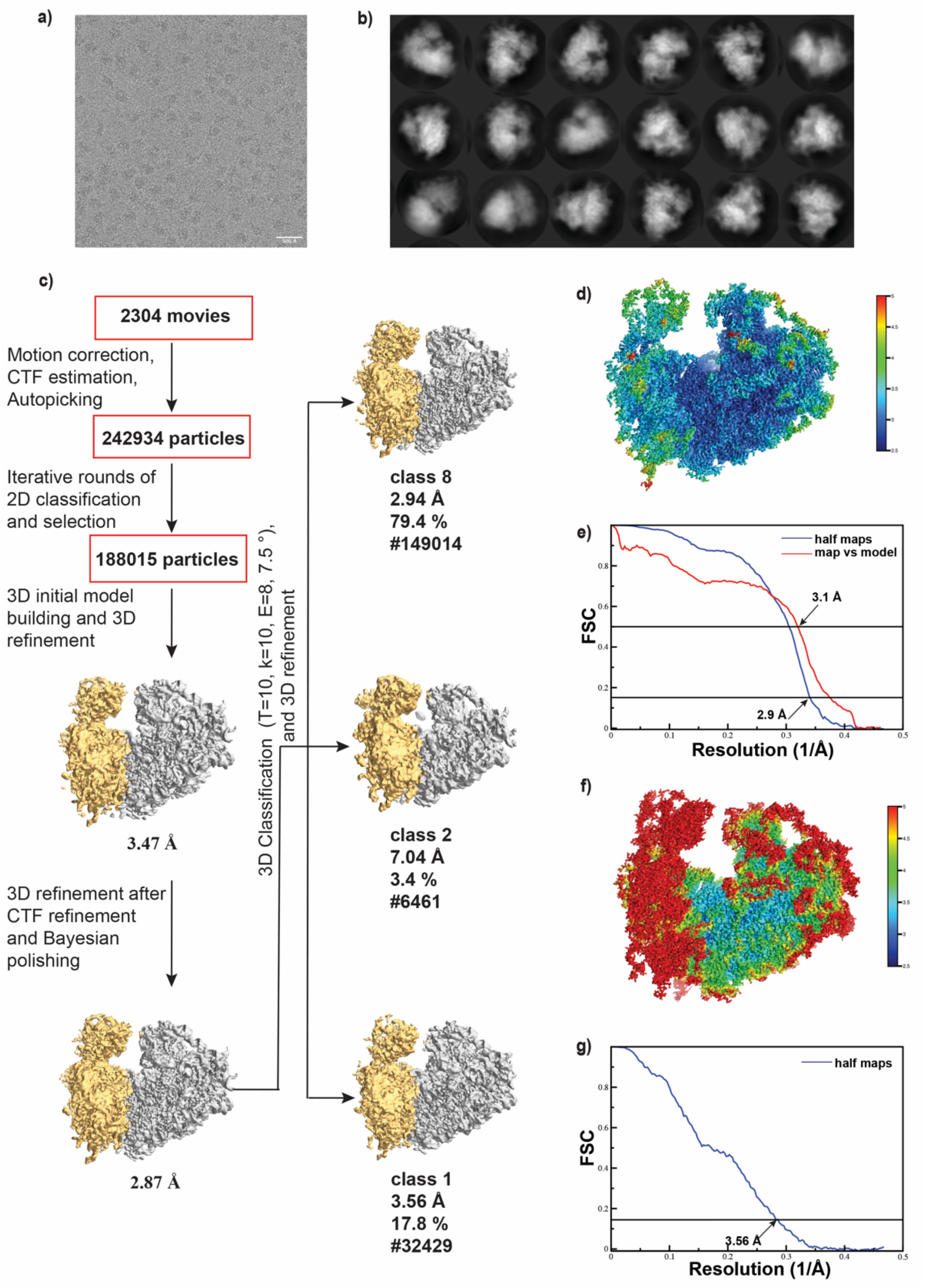
a)
b)
c)
2304 movies
Motion correction, CTF estimation, Autopicking
242934 particles
Iterative rounds of 2D classification and selection
188015 particles
3D initial model building and 3D refinement
3.47 Å
3D refinement after CTF refinement and Bayesian polishing
2.87 Å
3D Classification (T=10, k=10, E=8, 7.5 °), and 3D refinement
class 8
2.94 Å
79.4 %
#149014
class 2
7.04 Å
3.4 %
#6461
class 1
3.56 Å
17.8 %
#32429
d)
e)
half maps
map vs model
3.1 Å
2.9 Å
FSC
Resolution (1/Å)
f)
g)
half maps
3.56 Å
FSC
Resolution (1/Å)

h)

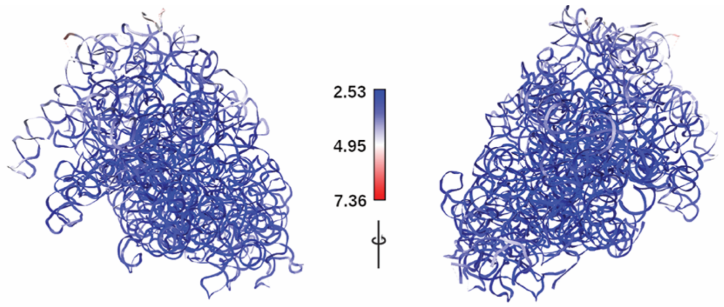


i)

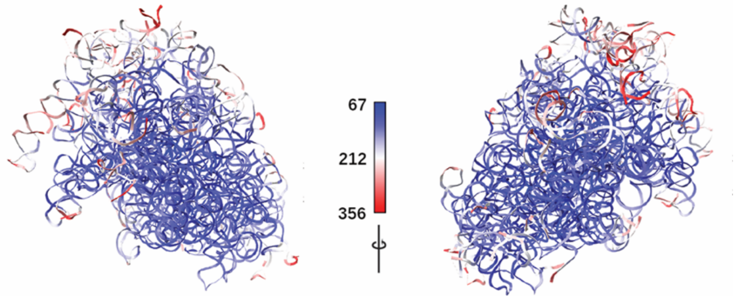


**Supplementary Figure 2. Cryo-EM data processing workflow for WT ribosomes, dataset 1**

**a)** A representative micrograph of WT ribosomes on carbon-coated holey grids showing well-distributed particles. **b)** 2D class averages of the 80S ribosomes of dataset 1 with the particles extracted using a box size of 360 pixels. **c)** Cryo-EM data processing workflow showing the steps followed to reach the final reconstruction using RELION. The maps shown here are the unsharpened maps and the resolutions indicated are after the post-processing step with a mask around the molecule. After 3D classification, 3 reasonable classes were obtained, and the largest class, class 8 with most particles resulted in highest resolution, class 1 was of intermediate resolution, which varied in the global 40S movement from class 8, and class 2 had very few particles and no reconstruction was performed. **d)** Local resolution of the final 3D reconstruction obtained from class 8. It is evident that resolution is not uniform across the map, and the blue coloured regions are at high resolution and the red coloured regions are at low resolution. Intrinsic flexibility of the 40S and the peripheral regions results in lower resolution. **e)** Fourier shell correlation (FSC) curve for the final 3D reconstruction of class 8. The gold-standard FSC calculated from the half-maps of the final refinement @ 0.143 is 2.9 Å and the map vs model FSC @ 0.5 is 3.1 Å. **f)** Local resolution of 3D reconstruction obtained from class 1 that has a global 40S movement. Due to high flexibility and 40S movement, the periphery and the 40S are at lower resolution than class 8. **g)** Fourier shell correlation (FSC) curve for the 3D reconstruction of class 1. The gold-standard FSC calculated from half-maps of the final refinement @ 0.143 is 3.6 Å. **h, i**) Two views of 28S rRNA model coloured according to the map values estimated with LocalRes, and with B-factor values after refinement. The range of the local resolution (Å), and the B-factor values (in Å$^2$) are shown.

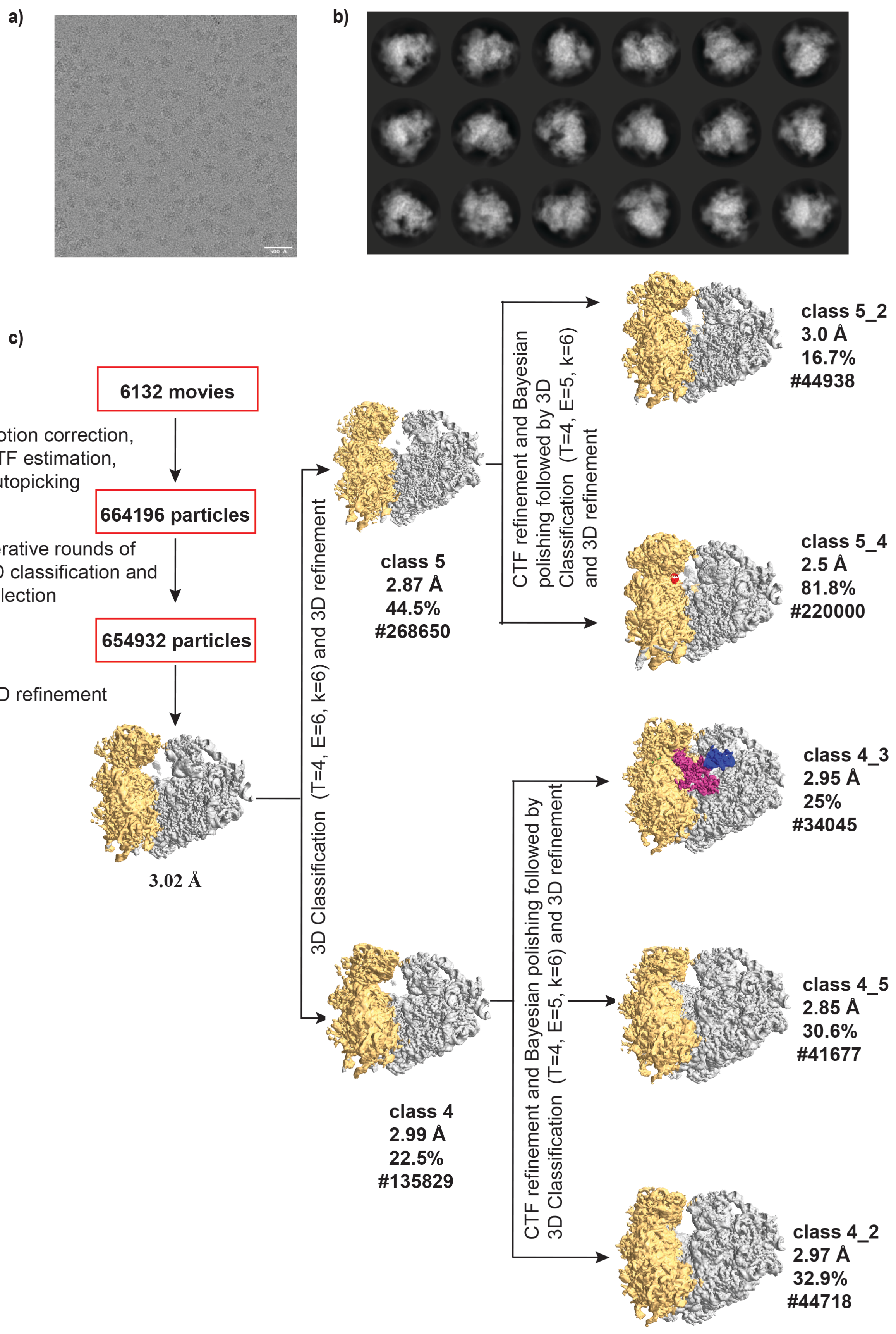
a)
b)
c)
6132 movies
Motion correction,
CTF estimation,
Autopicking
664196 particles
Iterative rounds of
2D classification and
selection
654932 particles
3D refinement
3.02 Å
3D Classification (T=4, E=6, k=6) and 3D refinement
class 5
2.87 Å
44.5%
#268650
class 4
2.99 Å
22.5%
#135829
CTF refinement and Bayesian polishing followed by 3D Classification (T=4, E=5, k=6) and 3D refinement
class 5_2
3.0 Å
16.7%
#44938
class 5_4
2.5 Å
81.8%
#220000
CTF refinement and Bayesian polishing followed by 3D Classification (T=4, E=5, k=6) and 3D refinement
class 4_3
2.95 Å
25%
#34045
class 4_5
2.85 Å
30.6%
#41677
class 4_2
2.97 Å
32.9%
#44718

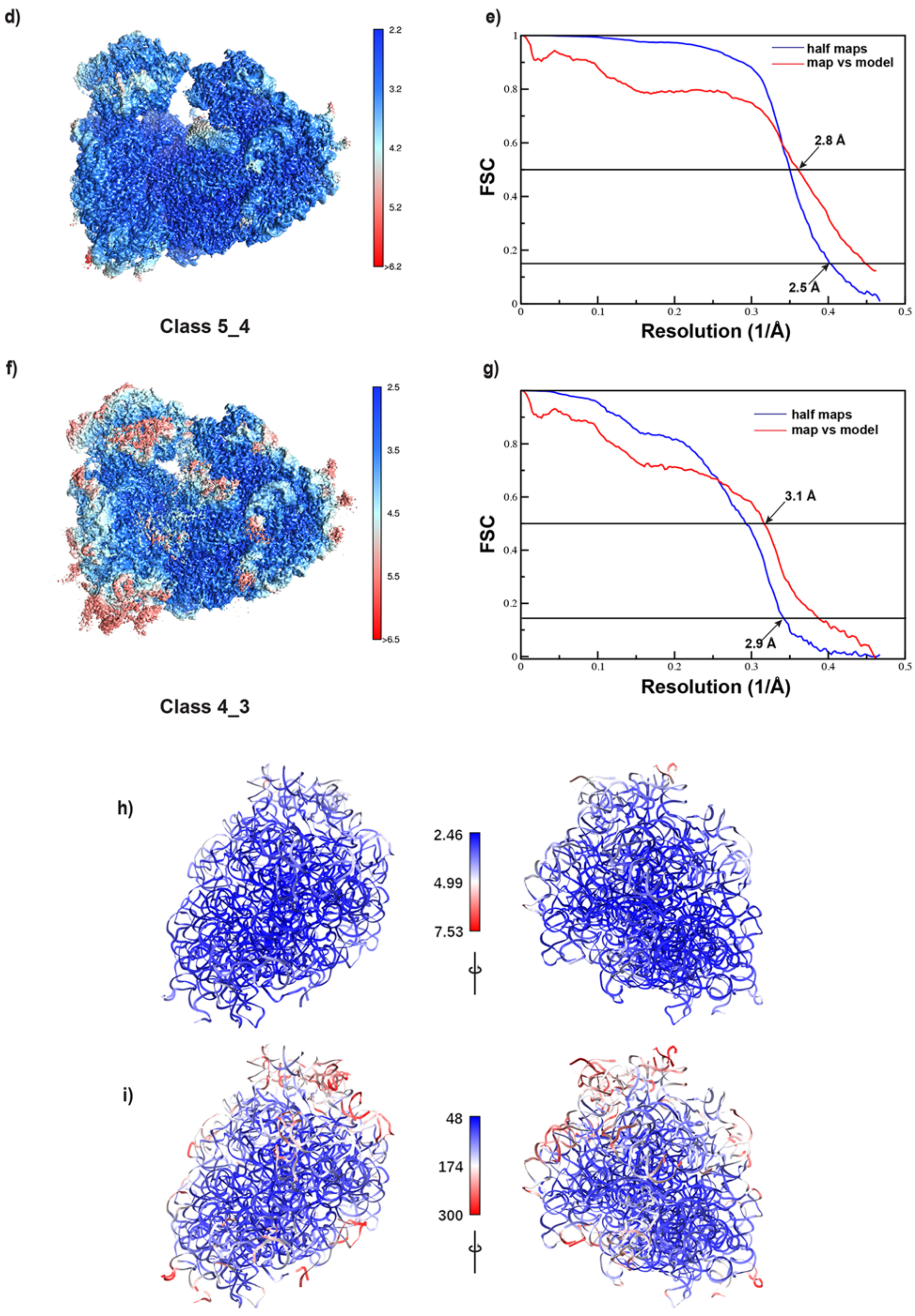


**Supplementary Figure 3. Cryo-EM data processing workflow for the FMR1 KO ribosomes, dataset 1**

**a)** Representative micrograph of FMR1 KO ribosomes on carbon-coated holey grids showing well-distributed particles. **b)** 2D class averages of the FMR1 KO ribosomes with a box size of 384 pixels. **c)** Cryo-EM data processing workflow showing the steps followed to reach the final reconstructions with RELION. After extensive 3D classification, multiple classes were obtained with different factors bound to them. Initial round gave two good classes (class 4 and class 5), which varied in 40S movement but when classified further, class 4 gave rise to three distinct populations (class 4_2, 4_3 and 4_5), and class 5 gave rise to two distinct populations (class 5_2 and class 5_4). Class 5_4 has the LYAR factor (coloured in red) bound, and class 5_2 differs in the 40S head movement from class 5_4, whereas the classes from class 4 had a mixture of empty and dormant ribosomes. Classes 4_2 and 4_5 were empty (no factors bound) and class 4_3 had eEF2, SERBP1, eIF5a and P0, and each factor is coloured differently with the eukaryotic elongation factor (eEF2) in magenta, SERBP1 in green, acidic protein P0 in blue, and eIF5a in forest green. All these maps are unsharpened maps with a threshold in the range of 0.03-0.07 and hide dust feature between 10-30. To colour the individual factors, colour zone option in ChimeraX was used with zone radii of 6-12. The resolutions of the reconstructions were calculated with the 80S mask. **d)** The local resolution plot of class 5_4 with LYAR bound. Much of the map is better resolved at higher resolution (2.2-3.5 Å). The 40S subunit is relatively more ordered in this class when compared to other maps. **e)** Fourier shell correlation (FSC) curve for the 3D reconstruction of class 5_4. The gold standard FSC calculated from the half-maps of the final refinement @ 0.143 is 2.5 Å, and the map vs model FSC @ 0.5 is 2.8 Å**. f)** The local resolution plot of class 4_3 with eEF2, SERBP1 and P0 bound. Much of the map is well-resolved at higher resolution (2.5-4 Å), but the peripheral regions and some parts of acidic protein P0 and eEF2 have lower resolution (>4 Å) due to intrinsic flexibility. **g)** Fourier shell correlation (FSC) curve for the 3D reconstruction of class 4_3. The gold standard FSC calculated from the half-maps of the final refinement @ 0.143 is 2.9 Å and the map vs model FSC @ 0.5 is 3.1 Å. Note that in the final deposited models, LYAR, eIF5a have not been included. **h, i**) Two views of 28S rRNA model (population with eEF2, class 4_3) coloured according to the map values estimated with LocalRes, and with B-factor values after refinement. The range of the local resolution (Å), and the B-factor values (in $Å^2$) are shown. The plots for class 5_4 are shown in Figure 4a & 4b.

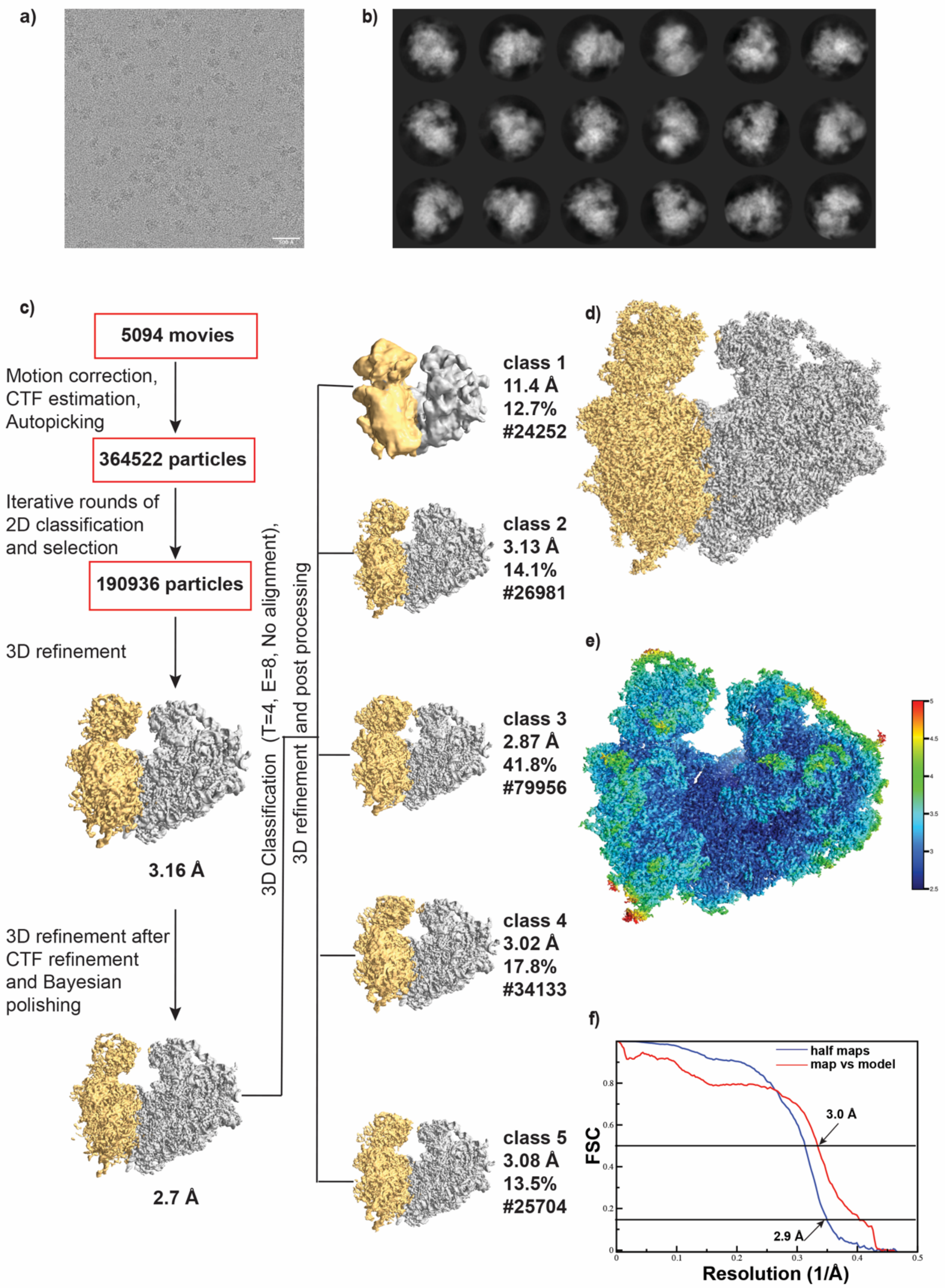
a)
b)
c)
5094 movies
Motion correction,
CTF estimation,
Autopicking
364522 particles
Iterative rounds of
2D classification
and selection
190936 particles
3D refinement
3.16 Å
3D refinement after
CTF refinement
and Bayesian
polishing
2.7 Å
3D Classification (T=4, E=8, No alignment),
3D refinement and post processing
class 1
11.4 Å
12.7%
#24252
class 2
3.13 Å
14.1%
#26981
class 3
2.87 Å
41.8%
#79956
class 4
3.02 Å
17.8%
#34133
class 5
3.08 Å
13.5%
#25704
d)
e)
f)
half maps
map vs model
3.0 Å
2.9 Å
FSC
Resolution (1/Å)
0
0.1
0.2
0.3
0.4
0.5
0.2
0.4
0.6
0.8

g)

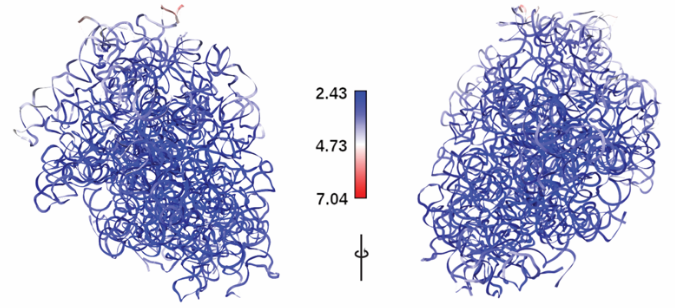


h)

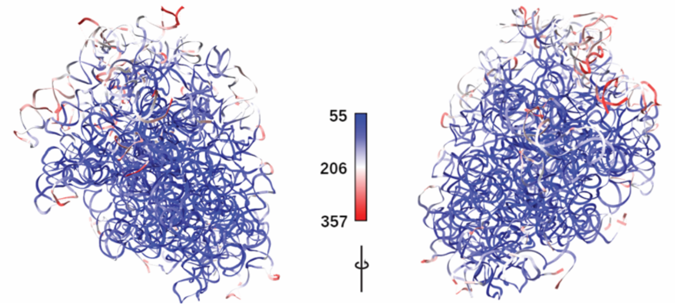


**Supplementary Figure 4. Cryo-EM data processing workflow for the WT ribosomes, dataset 2**

**a)** Representative micrograph of the WT ribosomes on carbon-coated holey grids showing well-distributed particles. **b)** 2D class averages are shown where different orientations are observed, and the box size of the classes is 384 pixels. **c)** Cryo-EM data processing workflow showing the steps followed to reach the final reconstruction with RELION. After Bayesian polishing, the reconstruction reached a resolution of 2.7 Å, which when classified, in 3D gave rise to five populations (class 1-5). Class 3 being the largest class, has the highest resolution, whereas classes 2, 4, and 5 had intermediate resolution, with class 1 being the lowest. The classes were likely separated based on the 40S movement, but additional factors were not found in any of the populations. Since class 3 was better resolved, it was used for the final reconstruction. **d)** The final reconstruction of class 3 obtained from this dataset. The large subunit has been coloured grey and the small subunit in gold. **e)** The local resolution map of the final reconstruction of class 3. It shows that most areas have very high resolution (2.5-4 Å). Only a few peripheral regions have low resolution (>4 Å) due to intrinsic flexibility. **f)** Fourier shell correlation (FSC) curve for the final reconstruction. The gold standard FSC calculated from half-maps of the final refinement @ 0.143 is 2.9 Å and the map vs model FSC @ 0.5 is 3 Å. **g, h**) Two views of 28S rRNA model coloured according to the map values estimated with LocalRes, and with B-factor after refinement. The range of the local resolution (Å), and the B-factor (in Å$^2$) are shown.

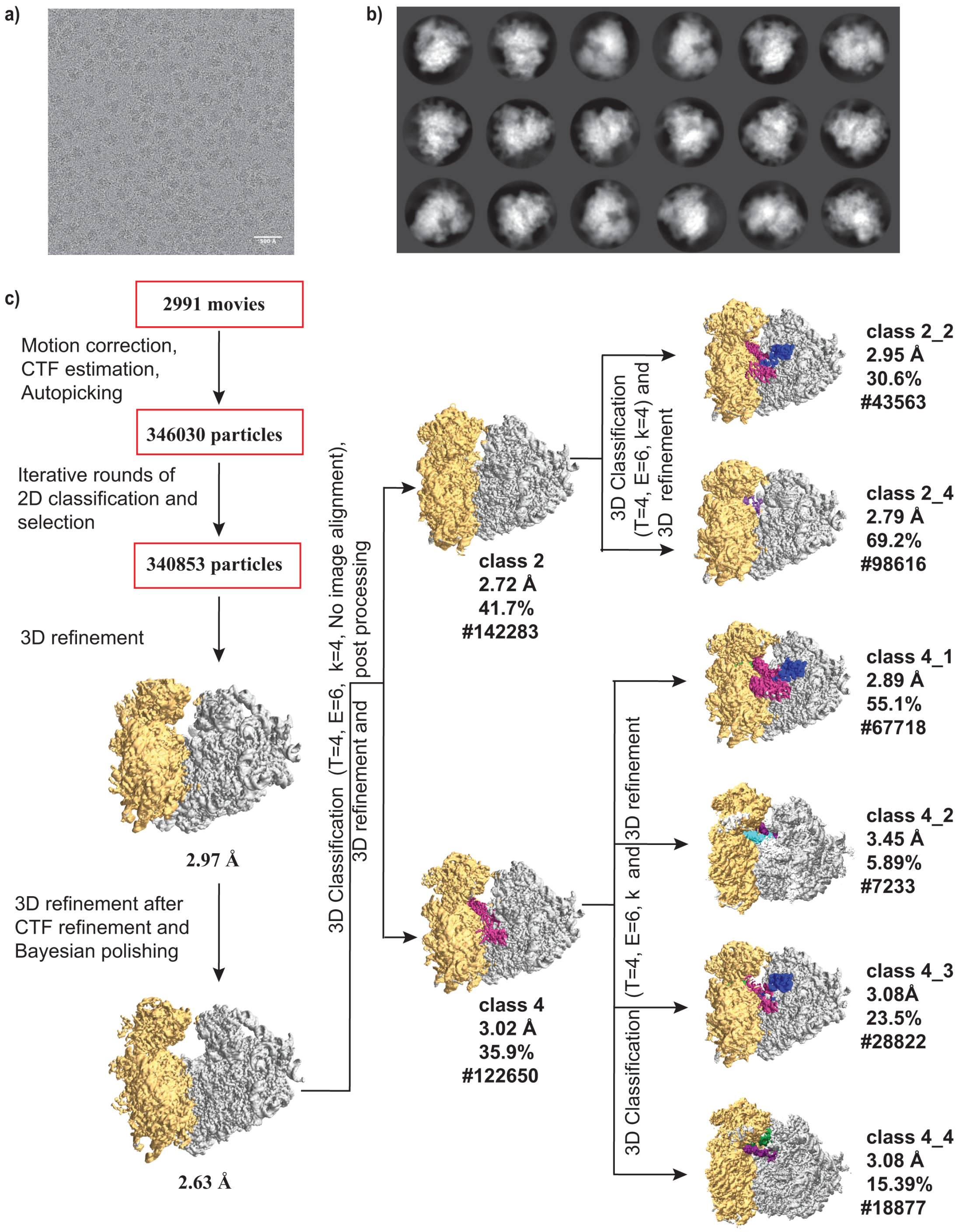
a)
b)
c)
2991 movies
Motion correction,
CTF estimation,
Autopicking
346030 particles
Iterative rounds of
2D classification and
selection
340853 particles
3D refinement
2.97 Å
3D refinement after
CTF refinement and
Bayesian polishing
2.63 Å
3D Classification (T=4, E=6, k=4, No image alignment),
3D refinement and post processing
class 2
2.72 Å
41.7%
#142283
class 4
3.02 Å
35.9%
#122650
3D Classification
(T=4, E=6, k=4) and
3D refinement
3D Classification (T=4, E=6, k and 3D refinement
class 2_2
2.95 Å
30.6%
#43563
class 2_4
2.79 Å
69.2%
#98616
class 4_1
2.89 Å
55.1%
#67718
class 4_2
3.45 Å
5.89%
#7233
class 4_3
3.08Å
23.5%
#28822
class 4_4
3.08 Å
15.39%
#18877

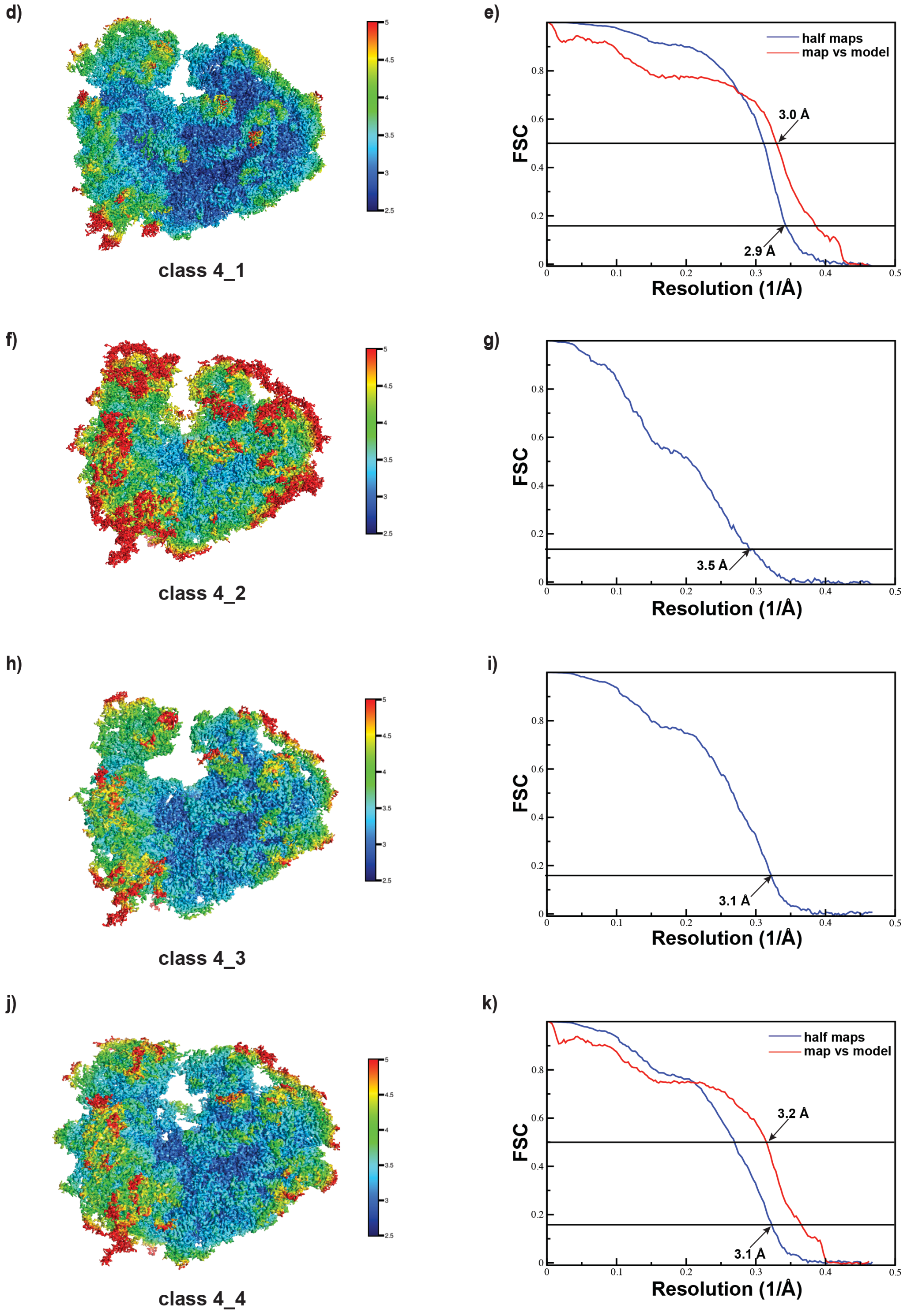
d)
class 4_1
e)
half maps
map vs model
3.0 Å
2.9 Å
FSC
Resolution (1/Å)
f)
class 4_2
g)
3.5 Å
FSC
Resolution (1/Å)
h)
class 4_3
i)
3.1 Å
FSC
Resolution (1/Å)
j)
class 4_4
k)
half maps
map vs model
3.2 Å
3.1 Å
FSC
Resolution (1/Å)

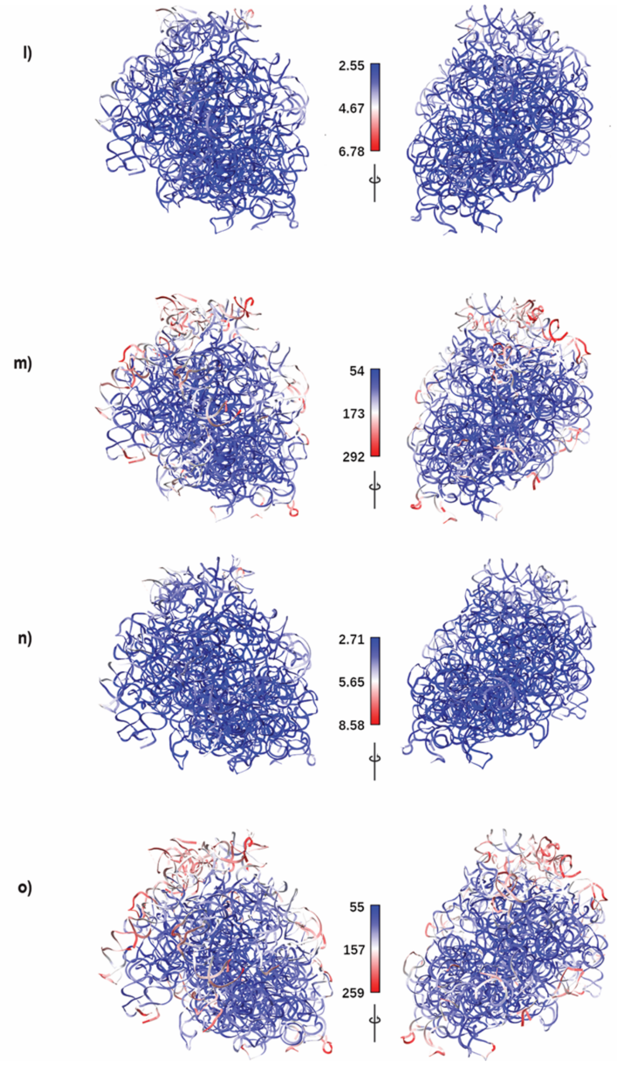


**Supplementary Figure 5. Cryo-EM data processing workflow for KO ribosomes, dataset 2**

**a)** Representative micrograph of the FMR1 KO ribosomes on carbon-coated grids showing abundant and well-distributed particles. **b)** 2D class averages are shown where different orientations are seen. The box size of the 2D classes is 384 pixels. **c)** Cryo-EM data processing workflow showing the steps followed in RELION. Highest resolution was obtained after polishing, following which extensive 3D classification was performed and multiple classes were obtained with different factors bound to them. In the first round, two classes (class 2 and class 4) indicated the presence of some additional density. Further classification was performed for both these classes with class 2 resulted in 2 populations (class 2_2 and class 2_4), while class 4 gave rise to 4 different populations. The factors bound in each class are coloured differently. Class 2_2 had the eukaryotic elongation factor (eEF2) (magenta), SERBP1 (green), and acidic protein P0 (blue), whereas, class 2_4 had P-site tRNA shown in purple (although at very low occupancy). On the other hand, two classes derived from class 4 (class 4_1 and class 4_3) had eEF2, SERBP1 and P0 and classes 4_2 and 4_4 had 2 tRNAs and an mRNA bound to them. Classes 4_1 and 4_3 differed in the 40S movement, whereas classes 4_2 and 4_4 varied in the tRNAs bound to them. Class 4_2 had A-site and P-site tRNAs and class 4_4 had P-site and E-site tRNAs respectively. The A-site, P-site and E-site tRNAs are coloured in cyan, purple and green respectively. All these maps are unsharpened maps with a threshold in the range of 0.022-0.03 and hide dust values of 10-30. To colour the individual factors, colour zone option in ChimeraX was used with zone radii of 6-12.

**d, f, h, j)** Local resolution maps of different sub-classes of class 4 with either eEF2, SERBP1 or tRNA bound. In all these classes, peripheral regions of the ribosomes are at lower resolution but the core is at higher resolution between 2.5-4 Å.

**e, g, i, k)** Fourier shell correlation (FSC) curves for different sub-classes of class 4 with gold standard FSC calculated between the two half-maps (blue) @ 0.143 and the map vs model (red) @ 0.5 (in class 4_1 and class 4_4).

**l, m**) Two views of 28S rRNA model (population with eEF2, class 4_1) coloured according to the map values estimated with LocalRes, and with B-factor. The range of the local resolution (Å), and the B-factor (in $Å^2$) are shown.

**n, o)** Two views of 28S rRNA model (population with tRNA, class 4_4) coloured according to the map values estimated with LocalRes, and with B-factor values. The range of the local resolution (Å), and the B-factor values (in $Å^2$) are shown.

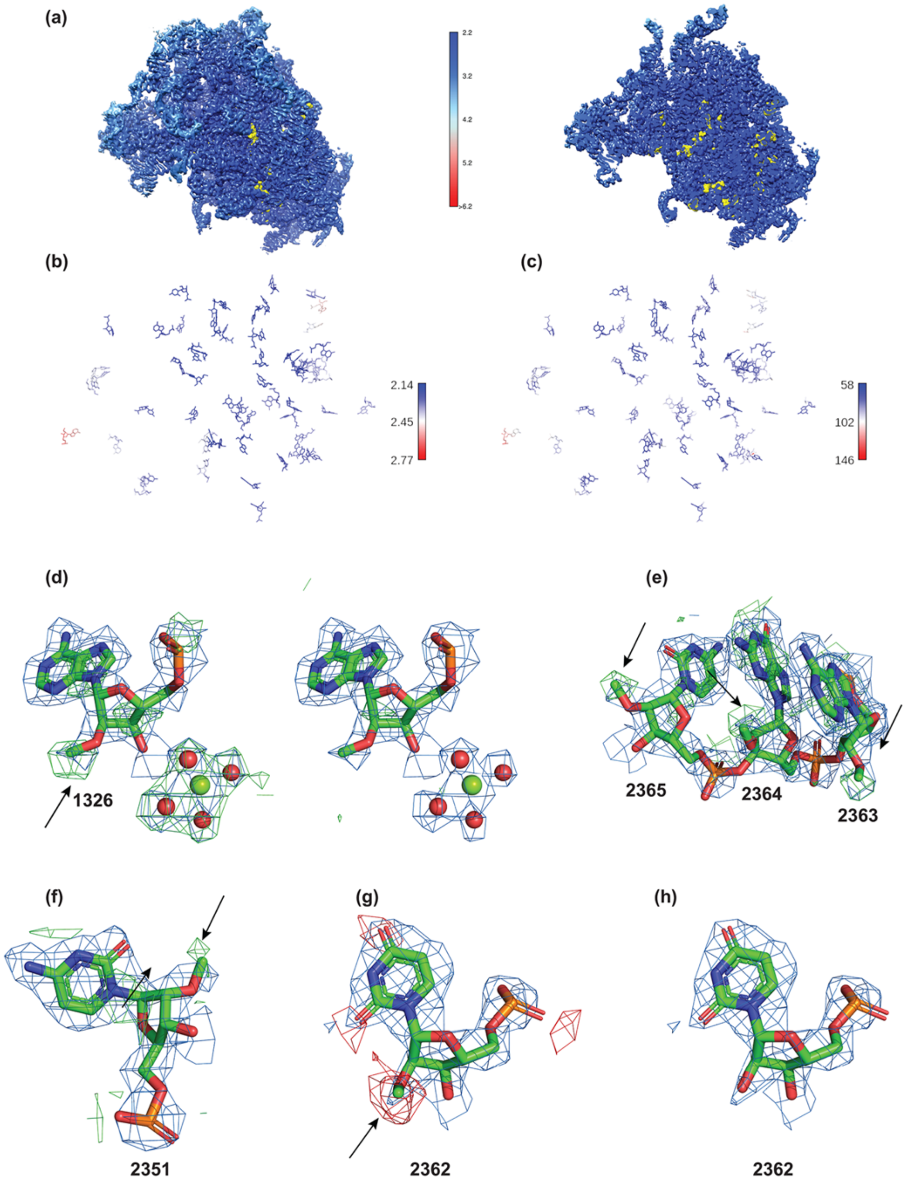


**Supplementary Figure 6. Local resolution plot of 28S rRNA and cryo-EM densities of select residues of 80S ribosome demonstrating the modelling of methyl groups on the ribose moiety.** FMR1 KO ribosome dataset 1(class 5_4), and the corresponding model was used to make this figure.

**a**) Local resolution plot of FMR1 KO ribosomes showing the cryo-EM map of only the 28S rRNA, and the modified residues (only the 2'-O-methylation from the model are shown in yellow spheres). The full map and the slice of the map (on the right side of the color key) are used to demonstrate the resolution of the interior of the rRNA, and to highlight the modified residues. The core of the 28S rRNA is better resolved with the periphery at lower resolution. Much of the modified residues in the current model are in the well-resolved region of the map. The local resolution of the complete map is shown in supplementary figure 3d.

**b & c**) For clarity, only the 2'-O methylated residues are shown. In panel b, these residues are colored according to local resolution (Å) with the color key showing the resolution range. In panel c, the same residues colored according to the final refined B-factor values (Å$^2$) are shown.

**d-h**) In all the panels, the final postprocess, auto-sharpen map (in blue) is shown around the atoms and the fofc map in green or red. (**d**) When residue 1326, is modelled as adenine (instead of A2M), a positive density (cryo-EM map, blue, 5 σ, and fofc map, green, 4 σ) is observed (here the modified residue is shown to highlight the positive density). The $Mg^{2+}$ and water molecules near this residue are also shown (cryo-EM map @ 3.5 σ ,and fofc @ 4 σ). After the addition of methyl group and ions/water and refinement, minimal difference density is observed around these residues (same sigma levels for the maps were used). (**e**) The three nucleotides stretch of 2363-2365, which are modified at 2'-O of the ribose are shown (cryo-EM map in blue and fofc map in green @ 4.5 σ). (**f**) Residue 2351, which also has 2'-O-methylation, but the difference map shows only a weak peak, perhaps due to lower resolution in this region (both maps @ 4.5 σ). (**g**) A methyl group was added to uracil 2362 and the model was refined. The difference map clearly shows a negative peak (colored red and @ 5 σ) indicating that the modelling of methyl group is not correct. The B-factor (ADP) values for the O2' for the model with methyl group is 81.3 $Å^2$ and without methyl group is 71.1 $Å^2$, indicating that the atom is most likely not present. (**h**) The density for the same residue with no methyl group is shown. The arrows in each panel denote the methyl group and the difference maps. We note that, such difference maps are useful, depending on the resolution, and one could also observe difference density (example, panel e, residue 2364 on the base), highlighting the limitations in the current calculations of such difference maps at intermediate resolutions.

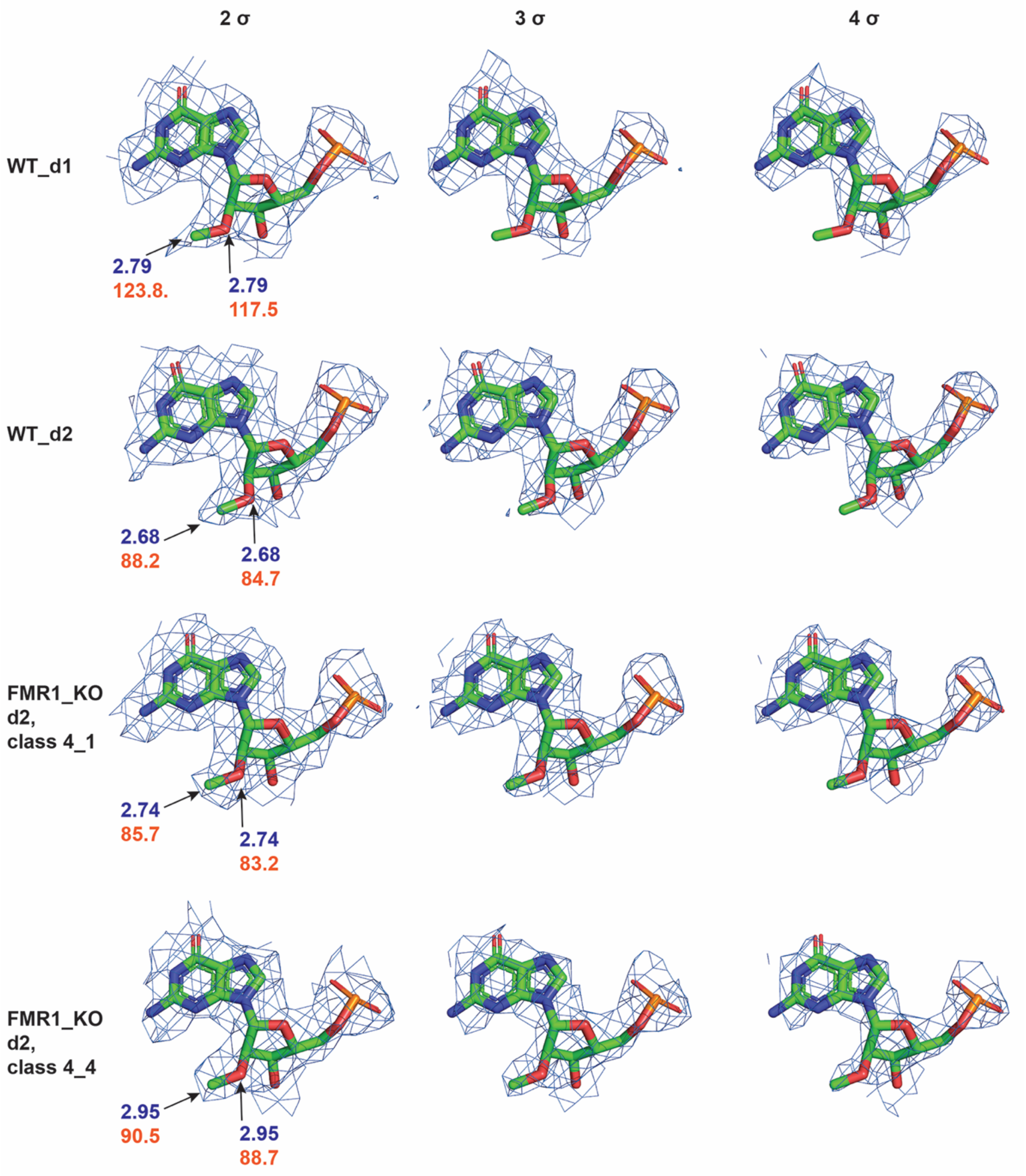


**Supplementary Figure 7. Density for methyl group of a selected residue across different ribosomal classes at different threshold values**

Cryo-EM density for residue 4618 in the two WT data sets and the two populations of FMR1-KO data sets. The modified residue is 2'-O-methyl guanine, which on visual inspection at high contour suggests a lack of methyl group in the WT datasets, but gives reasonable Q-score when modelled (supplementary table 2). In this figure, maps have been carved (2 Å) around the residue at different sigma levels (in PyMOL). While the FMR1 KO populations show density for the methyl group at higher contour levels, the corresponding density in the WT datasets is less prominent. In addition to the density at different contour levels, the local resolution (blue) and B-factor values for the O2' and methyl group of the ribose sugars for each dataset is given.

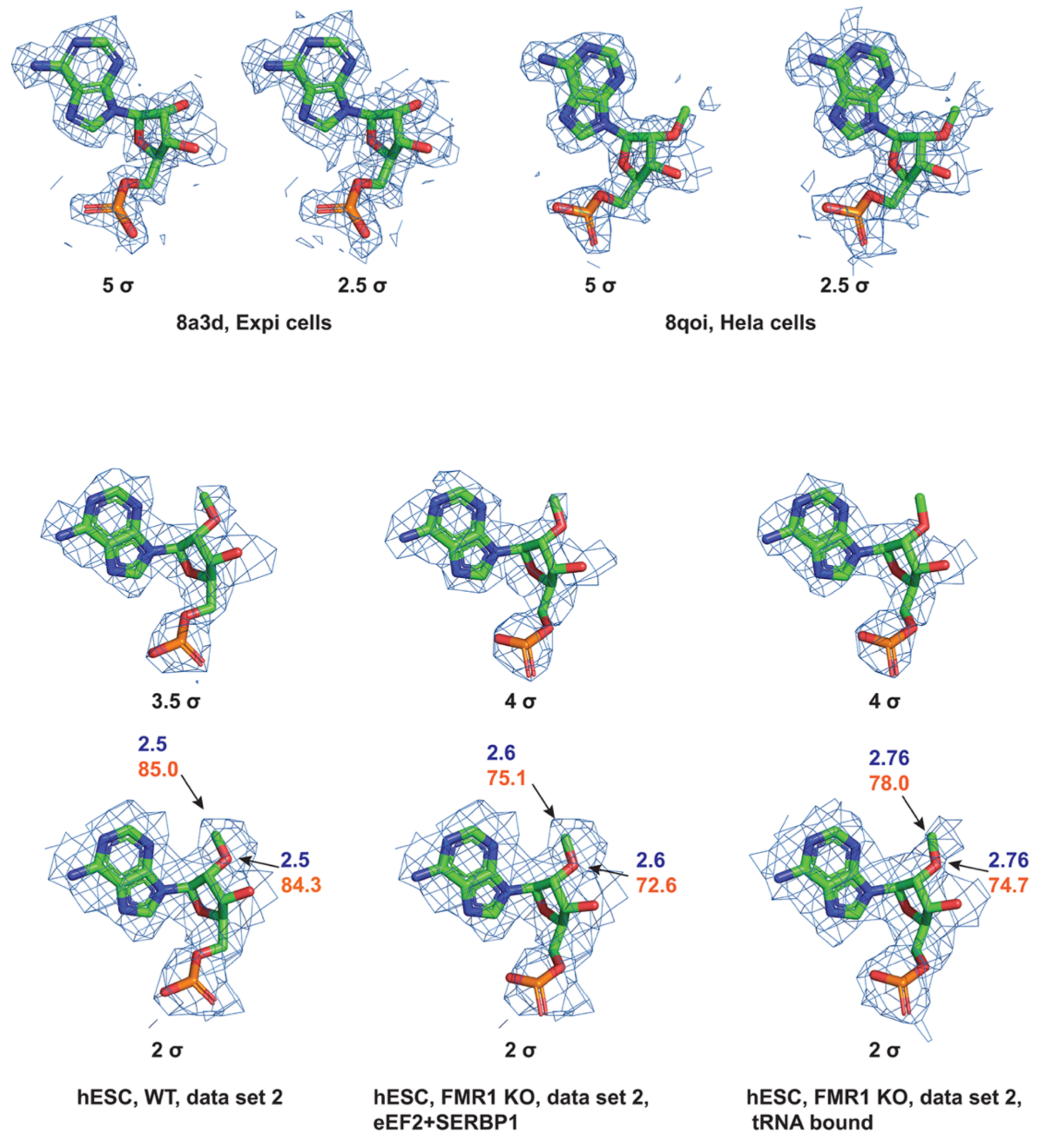


**Supplementary Figure 8. Analysis of methylation of residue 1323 in 28S rRNA in ribosome structures across human cell lines.**
Density for residue 1323 in 28S rRNA from multiple ribosome maps shown at two different contour levels. These include Expi cells (PDB-8a3d and EMD-15113), HeLa cells (PDB-8qoi and EMD-18539), WT and FMR1 KO cells from dataset 2. The resolutions of the Expi and HeLa datasets are very high (1.7 and 1.9 Å respectively), and clear differences in the density corresponding to 2'-O methyl group can be seen. In the hESC dataset 2, the density for the methyl group is clear in WT but no clear density is present in FMR1 KO tRNA-bound population, while residual density is present in the dormant ribosomal population. The resolutions and numbers of particles are different across these datasets, which might also affect the quality of the density. In fact, at a lower contour level, density for the methyl group can be observed in the tRNA-bound class. In addition to the density at different contour levels, the local resolution (blue) and B-factor values for the O2' atom and the methyl group of the ribose sugars is given for the datasets described in the manuscript. We also note that for another residue, 1881, the methyl group is not present in 8a3d model but is modelled in the current models and in 8qoi.

**Supplementary Table 1: Density for 2'-O-methylation in maps of different populations**

| Residues | WT dataset 1 class 8 | FMR1 KO dataset 1 class 4_3 | FMR1 KO dataset 1 class 5_4 | WT dataset 2 class 3 | FMR1 KO dataset 2 class 4_1 | FMR1 KO dataset 2 class 4_4 |
|---|---|---|---|---|---|---|
| 398-A2M | + | + | + | + (High) | + | -/? |
| 400-A2M | + | + | + | + (High) | + | + |
| 1316-OMG | + | + (High) | + | + | + | + |
| 1323-A2M | + (High) | + | + (High) | + (High) | + | - |
| 1326-A2M | + | + | + | + | + | + |
| 1340-OMC | + | + | + | + | + | + (High)/? |
| 1522-OMG | + | + | + | + | + | + |
| 1524-A2M | + | + | + | + (High)/? | + | + |
| 1534-A2M | + | + | + | + | + | + |
| 1625-OMG | + | + (High) | + | + | + (High)/? | + |
| 1871-A2M | + | + | + | + | + | + |
| 1881-OMC | + | + (High) | + | ? | + (High) | + (High) |
| 2351-OMC | + | + | + | + (High) | + (High) | + (High) |
| 2363-A2M | + | + | + | + | + | + |
| 2364-OMG | + | + | + (High) | + | + | + |
| 2365-OMC | + (High) | + | + | + (High) | + | + |
| 2401-A2M | + | + | + | + (High) | + | + |
| 2422-OMC | + | + | + | + | + | + |
| 2424-OMG | + (High) | + | + | + (High) | + (High) | + (High) |
| 2787-A2M | + | + | + (High) | + (High) | + | + |
| 2804-OMC | + | + | + | + | + | + |
| 2815-A2M | + (High) | + (High) | + | + (High) | + (High) | + |
| 2824-OMC | + | + | + | ? | + | + (High) |
| 2837-OMU | + (High) | + (High) | + | + (High) | + | -/? |
| 2861-OMC | + | + | + | + (High) | + | + |
| 2876-OMG | + (High) | + (High) | + | + (High) | + (High) | + |
| 3627-OMG | + | + | + | + | + | + (High) |
| 3701-OMC | + | + (High) | + | + | + (High) | + |
| 3718-A2M | -/ very high contour? | + | + | + | + | + (High)/? |
| 3724-A2M | Poor density | Poor density | + (High) | Poor density | Poor density | Poor density |
| 3744-OMG | + (High) | + (High) | + (High) | + | + | -/? |
| 3785-A2M | + | + | + | + | + | + |
| 3792-OMG | + (High) | + | + | + | + | + |
| 3808-OMC | + | + | + | + | + | + (High) |
| 3825-A2M | + | + | + | + | + | + |
| 3830-A2M | + | + | + | + (High) | + | + |
| 3841-OMC | + | + | + | + | + | + |
| 3867-A2M | + (High) | + (High) | + (High) | + | + (High) | -/? |
| 3869-OMC | + | + | + | + | + | + |
| 3887-OMC | + | + | + | + | + | + |
| 3899-OMG | + | + | + | + | + | + |

| | | | | | | |
|---|---|---|---|---|---|---|
| 3925-OMU | + | + | + | + | + | + (High) |
| 4196-OMG | + | + | + (High) | + | + | + |
| 4227-OMU | -/? | + (High) | + (High) | + (High) | + (High) | + (High) |
| 4228-OMG | + | + | + | + | + | + |
| 4306-OMU | + | + (High) | + | + | + | + (High) |
| 4370-OMG | + | +/? | + | + (High)/? | + | + |
| 4392-OMG | + (High) | + | + | + (High)/? | + | + |
| 4456-OMC | + | + | + | + | + (High) | + |
| 4494-OMG | + | + | + | + | + | + (High)/? |
| 4498-OMU | + (High) | + (High) | + (High) | + (High)/? | -/? | + |
| 4499-OMG | + | + | + | + (High) | + | + (High) |
| 4523-A2M | + | + | + | + | + | + |
| 4536-OMC | + (High) | + | + | + (High) | + (High) | + |
| 4571-A2M | + (High/poor density) | + (High/poor density) | + (High/poor density) | + (High) | + (High)/? | + |
| 4590-A2M | + (High) | - | + (High/not modelled) | + (High)/? | -/? | -/? |
| 4618-OMG | - | + (High) | + | - | + | + |
| 4620-OMU | -/? | + | + | + (High) | + | + (High) |
| 4623-OMG | + (High) | + | + | + | + | + |
| 4637-OMG | + (High) | + | + | + (High) | + (High) | + (High) |

| | |
|---|---|
| + | Present |
| - | Absent |
| ? | Ambiguous |
| + (High) | Present at high contour |

**Supplementary Table 2: MapQ score for 2' O-methyl groups (using Chimera and sharpened maps from the postprocess step, @ 0.6 sigma)**

| Residue[a] Number and modification | WT, d1, class 8 | FMR1 KO, d1, class 4_3 | FMR1 KO, d1, LYAR class 5_4 | WT, d2 class 3 | FMR1 KO, d2, eEF2 class 4_1 | FMR1 KO, d2, tRNA class 4_4 |
|---|---|---|---|---|---|---|
| 398, A2M | 0.83 | 0.81 | 0.89 | 0.84 | 0.69 | 0.61 |
| 400, A2M | 0.85 | 0.67 | 0.84 | 0.86 | 0.79 | 0.79 |
| 1316, OMG | 0.88 | 0.74 | 0.82 | 0.83 | 0.89 | 0.88 |
| 1323, A2M | 0.72 | 0.86 | 0.79 | 0.86 | 0.87 | **0.48** |
| 1326, A2M | 0.8 | 0.76 | 0.86 | 0.85 | 0.8 | 0.72 |
| 1340, OMC | 0.9 | 0.81 | 0.88 | 0.79 | 0.89 | 0.68 |
| 1522, OMG | 0.88 | 0.89 | 0.87 | 0.84 | 0.79 | 0.8 |
| 1524, A2M | 0.86 | 0.87 | 0.93 | 0.81 | 0.88 | 0.91 |
| 1534, A2M | 0.86 | 0.93 | 0.94 | 0.83 | 0.85 | 0.86 |
| 1625, OMG | 0.78 | 0.85 | 0.9 | 0.86 | 0.78 | 0.79 |
| 1871, A2M | 0.84 | 0.86 | 0.89 | 0.89 | 0.83 | 0.74 |
| 1881, OMC | 0.63 | 0.91 | 0.86 | 0.71 | 0.79 | 0.66 |
| 2351, OMC | 0.81 | 0.79 | 0.76 | 0.79 | 0.88 | 0.77 |
| 2363, A2M | 0.84 | 0.9 | 0.88 | 0.87 | 0.86 | 0.86 |
| 2364, OMG | 0.77 | 0.85 | 0.87 | 0.8 | 0.86 | 0.86 |
| 2365, OMC | 0.74 | 0.8 | 0.88 | 0.81 | 0.81 | 0.82 |
| 2401, A2M | 0.75 | 0.85 | 0.84 | 0.82 | 0.87 | 0.74 |
| 2422, OMC | 0.68 | 0.71 | 0.93 | 0.85 | 0.89 | 0.73 |
| 2424, OMG | 0.78 | 0.84 | 0.8 | 0.75 | 0.85 | 0.89 |
| 2787, A2M | 0.87 | 0.81 | 0.8 | 0.83 | 0.8 | 0.74 |
| 2804, OMC | 0.87 | 0.88 | 0.91 | 0.84 | 0.88 | 0.76 |
| 2815, A2M | 0.55 | 0.7 | 0.88 | 0.79 | 0.85 | 0.85 |
| 2824, OMC | 0.84 | 0.71 | 0.9 | 0.78 | 0.83 | 0.86 |
| 2837, OMU | 0.82 | 0.81 | 0.84 | 0.85 | 0.85 | 0.86 |
| 2861, OMC | 0.93 | 0.65 | 0.73 | 0.83 | 0.93 | 0.78 |
| 2876, OMG | 0.85 | **0.19** | 0.82 | 0.81 | 0.8 | 0.8 |
| 3627, OMG | 0.91 | 0.76 | 0.9 | 0.83 | 0.85 | 0.6 |
| 3701, OMC | 0.77 | 0.6 | 0.72 | 0.74 | 0.89 | 0.8 |
| 3718, A2M | 0.58 | 0.79 | 0.83 | 0.89 | 0.74 | 0.75 |
| 3724, A2M | 0.54 | 0.57 | 0.71 | 0.79 | 0.73 | 0.8 |
| 3744, OMG | 0.79 | 0.61 | 0.85 | 0.83 | 0.89 | 0.84 |
| 3785, A2M | 0.84 | 0.88 | 0.92 | 0.88 | 0.91 | 0.87 |
| 3792, OMG | 0.77 | 0.66 | 0.9 | 0.89 | 0.9 | 0.89 |
| 3808, OMC | 0.91 | 0.84 | 0.84 | 0.77 | 0.79 | 0.72 |
| 3825, A2M | 0.91 | 0.92 | 0.85 | 0.87 | 0.89 | 0.86 |
| 3830, A2M | 0.71 | 0.74 | 0.89 | 0.78 | 0.67 | 0.84 |
| 3841, OMC | 0.93 | 0.84 | 0.91 | 0.87 | 0.9 | 0.85 |

| | | | | | | |
|---|---|---|---|---|---|---|
| 3867, A2M | 0.72 | 0.6 | 0.7 | 0.7 | 0.76 | **0.3** |
| 3869, OMC | 0.8 | 0.91 | 0.77 | 0.81 | 0.89 | 0.83 |
| 3887, OMC | 0.85 | 0.91 | 0.91 | 0.8 | 0.89 | 0.79 |
| 3899, OMG | 0.87 | 0.82 | 0.82 | 0.88 | 0.85 | 0.81 |
| 3925, OMU | 0.83 | 0.86 | 0.9 | 0.9 | 0.86 | 0.74 |
| 4196, OMG | 0.78 | 0.9 | 0.93 | 0.8 | 0.83 | 0.82 |
| 4227, OMU | 0.43 | 0.49 | 0.87 | 0.63 | 0.75 | 0.62 |
| 4228, OMG | 0.79 | 0.88 | 0.8 | 0.79 | 0.78 | 0.8 |
| 4306, OMU | 0.79 | 0.82 | 0.93 | 0.87 | 0.84 | 0.75 |
| 4370, OMG | 0.76 | 0.74 | 0.66 | 0.8 | 0.74 | 0.82 |
| 4392, OMG | 0.79 | 0.89 | 0.88 | 0.76 | 0.87 | 0.83 |
| 4456, OMC | 0.83 | 0.87 | 0.88 | 0.82 | 0.82 | 0.8 |
| 4494, OMG | 0.8 | 0.93 | 0.9 | 0.9 | 0.81 | 0.76 |
| 4498, OMU | 0.68 | 0.83 | 0.73 | 0.86 | 0.82 | 0.81 |
| 4499, OMG | 0.81 | 0.79 | 0.87 | 0.91 | 0.88 | 0.9 |
| 4523, A2M | 0.86 | 0.88 | 0.83 | 0.82 | 0.87 | 0.88 |
| 4536, OMC | 0.84 | 0.87 | 0.91 | 0.81 | 0.84 | 0.86 |
| 4571, A2M | 0.83 | 0.76 | 0.86 | 0.88 | 0.76 | 0.76 |
| 4590, A2M[b] | - | - | - | - | - | - |
| 4618, OMG[c] | 0.6 | 0.75 | 0.91 | 0.76 | 0.8 | 0.81 |
| 4620, OMU | 0.77 | 0.82 | 0.86 | 0.84 | 0.82 | 0.75 |
| 4623, OMG | 0.81 | 0.74 | 0.92 | 0.93 | 0.86 | 0.83 |
| 4637, OMG | 0.83 | 0.82 | 0.87 | 0.86 | 0.78 | 0.82 |

a- Methylation in residues 1323, 3627, 3718 are modelled from cryo-EM maps, but they are not observed in RiboMethSeq (Ninochka D'Souza et al 2025). Residues including 1760, 3760, 3818, 3944, 4042, 4054, 4590 have not been modelled in any of the current models, as some are in periphery with no density or unclear density, but observed in RiboMethSeq data. In other PDBs, 8a3d, 8qoi and 8qyx, clear density for 3627 and 3718 is observed, and also 3818 has been modelled. In 8a3d, residues 1323 and 1881 are not modelled with methylation.

b- Modification in residue 4590 is present in PDBs 8acd and 8qoi but not in 8qyx. In the current data sets, it is unclear if there is extra density for methyl group, though in WT dataset 1 there is some density but not in other data sets. Hence, not modelled.

c- The modification on residue 4618 is not modelled in the WT data sets. However, OMG was modelled, refined and Q score was calculated for both the WT data sets, which show respectable scores though the density is poor visually (Supplementary Table 1). Similarly, the values highlighted in green (for KO datasets) correspond to cases in which the methyl group is modelled but shows poor density and lower Q-scores.
